# Dendritic spines implement specific connectivity and support neuronal ensembles in cortical circuits

**DOI:** 10.64898/2026.06.07.730704

**Authors:** Dean Geckt, Netanel Ofer, Michael W. Reimann, Rafael Yuste, Idan Segev

## Abstract

Dendritic spines are the principal postsynaptic targets of excitatory synapses, yet the network logic governing their organization and contribution to cortical computation remains unclear. Here, we combine the MICrONS ultrastructural reconstruction of mouse visual cortex with matched *in vivo* two-photon calcium imaging to explore the network roles of dendritic spines. Structurally, we show that spine density predicts the number and diversity of both excitatory and inhibitory presynaptic partners, supporting the long-standing “connectivity and diversity” hypothesis. Beyond expanding input space, spines exhibit distinct network-level principles: excitatory axons preferentially target spines with increasing axonal distance from the soma; highly broadcasting (“hub”) neurons preferentially innervate spines; and neurons sharing presynaptic partners preferentially route their own outputs onto spines, revealing an input-to-spine/output-to-spine wiring correlations. Functionally, we find that these same structural motifs are found in neuronal ensembles, defined as groups of neurons with correlated calcium activity. Ensemble members are interconnected at more than twice the rate of spatially matched controls; these synapses connecting them are routed almost exclusively onto dendritic spines; and their shared presynaptic partners are almost exclusively inhibitory neurons that themselves also preferentially target spines. We suggest that local recurrent spine-targeted excitation effectively binds ensemble members into a coactive subnetwork, while shared spine-directed inhibition gates, synchronizes, and stabilizes their collective activity, providing a circuit-level architecture that may support attractor dynamics, pattern completion, and other cortical computations.

## Introduction

Dendritic spines are among the most distinctive features of cortical excitatory neurons and constitute the principal postsynaptic targets of excitatory synapses. For more than a century, they have been proposed to expand neuronal connectivity by increasing the number and diversity of synaptic inputs - the “connectivity and diversity” hypothesis (Yuste, 2011). Although the cellular properties of dendritic spines have been extensively characterized, their organizing principles at the scale of cortical circuits remain largely unexplored. In particular, it is unclear whether spines merely increase the number of synaptic contacts available to a neuron or whether they may implement broader network-level wiring rules that shape cortical computations.

At the subcellular level, synaptic targeting follows well-established biases: excitatory synapses preferentially innervate dendritic spines, whereas inhibitory synapses primarily target dendritic shafts (Kwon et al., 2019; Megías et al., 2001; Peters et al., 1991). Nevertheless, the shaft is not exclusively an inhibitory domain; a substantial fraction of excitatory synapses also contacts dendritic shafts directly and may constitute a more structurally flexible synaptic population (Fanutza et al., 2025). These compartment-specific observations, however, provide limited insight into how spines contribute to the organization of cortical connectivity. Indeed, cortical connectivity has long been recognized as non-random, exhibiting long-tailed degree distributions (Piazza et al., 2025; Brunel, 2016), common-neighbor motifs (Perin et al., 2011), assortative wiring (Ko et al., 2011; Cossell et al., 2015), and rich-club organization (Van Den Heuvel and Sporns, 2011). However, whether dendritic spines participate in establishing these higher-order principles remains unknown.

Ultimately, these structural questions are inseparable from functional questions. Cortical computation is widely thought to emerge from neuronal ensembles, groups of neurons that are coactive during sensory stimulation or behavior and also present in spontaneous activity (Yuste, 2026). Since Donald Hebb, such ensembles (i.e. assemblies) have often been assumed to arise from strengthened recurrent excitatory connectivity. Yet recent studies combining functional measurements with circuit anatomy have challenged this simple view: in mouse visual cortex, ensemble membership is not accompanied by stronger or more probable monosynaptic excitatory connections (Wagner-Carena et al., 2025), and experimentally induced ensembles can emerge without measurable changes in synaptic strength (Alejandre-García et al., 2022). In modeling studies (Ecker et al., 2024), it has been found that extrinsic input is a better predictor of ensemble membership than intrinsic connectivity, but intrinsic connectivity becomes more informative if membership in connectivity motifs is taken into account. However, these simulations have not yet been confirmed experimental, hence the structural mechanisms that generate a functional ensemble remain unresolved.

Here, we address these questions by combining the MICrONS large-scale electron microscopy reconstruction of the mouse visual cortex, comprising approximately 337 million synapses, with matched *in vivo* two-photon calcium imaging of a subpopulation of neurons whose internal connectivity is also available (MICrONS Consortium, 2025). Our primary connectivity analyses used a manually proofread local micro-column (Schneider-Mizell et al., 2025), allowing reliable identification of synaptic partners while retaining access to the broader volumetric circuit. We analyze these data to determine how spine density and spine targeting relate to synaptic diversity, axonal broadcasting, shared-input architecture, and functional ensemble membership. Our findings reveal that dendritic spines do not only expand synaptic connectivity, but that they support coordinated neuronal activity.

## Results

### Topological analysis at two different spatial scales

Our analysis evaluates this dataset across two distinct topological scales. The first is a densely reconstructed local micro-column (Schneider-Mizell et al., 2025), containing ∼146,000 intrinsic synapses exclusively connecting a highly curated subset of 1,298 fully reconstructed and manually identified pre- and post-synaptic neurons within the subcircuit (filtered from an initial set of 1,352; see Methods). This micro-column is situated at the center of the VISp region, spanning a 100 × 100 µm lateral area across the entire cortical depth. The second scale is a large-scale ‘global’ network that captures the extensive connectivity between the micro-column and the surrounding cortical tissue, comprising ∼4.6 million incoming synapses projecting onto the micro-column from the broader 1 mm^3^ volume (and outside of it), alongside ∼1 million outgoing synapses originated from the micro-column and projecting to the rest of the 1 mm^3^ volume. For all these synaptic connections, a prediction of its ultrastructural target - spine or shaft - was estimated to reach 95% accuracy (Pedigo et al., 2026).

The local micro-column represents a uniquely high-fidelity subset of this volume because it is a complete axon-proofread dataset. While automated segmentation algorithms can reconstruct large-scale volumes, they remain prone to “splits” (fragmented processes) and “mergers” (incorrectly joined processes). To mitigate these errors, MICrONS utilized human annotators to review and correct the axonal arbors within this subcircuit. This manual verification is particularly important for axons, which are significantly thinner and more morphologically complex than dendrites, making them highly susceptible to automated errors. By only utilizing an axon-proofread dataset, we ensure that the identified synaptic partnerships accurately reflect biological ground truth rather than segmentation artifacts, providing a reliable foundation for our analysis. Furthermore, despite its localized scale, this verified subcircuit was shown to serve as a faithful representative of the broader VISp region in both its cell type composition and their relative proportions (Zhang et al., 2026; Schneider-Mizell et al., 2025).

The distinction between these two topological scales is governed by specific analytical “rules” that ensure biological validity across varying levels of reconstruction detail. The local micro-column serves as our primary foundation for connectivity analysis; because it is axon-proofread, we can construct a high-confidence adjacency matrix where every synapse is traced back through its verified axonal arbor to its specific pre-synaptic origin. However, to address questions requiring greater statistical scale, we utilize the ‘global’ network under two conditions.

First, when analyzing total synaptic input richness (e.g., Fig. 1), we leverage the ∼4.6 million incoming synapses. This is justified by the relative morphological stability of dendrites - which are less prone to automated segmentation errors than axons - and the fact that this analysis requires only the accurate detection of a synapse on a known post-synaptic target, rather than the identity of the pre-synaptic partner. Second, when quantifying axonal output (e.g., Fig. 3), we utilize the ∼1 million outgoing synapses. Here, we rely on the proofread axons within the micro-column to calculate total synaptic release, a metric that remains robust regardless of the specific identities of the target neurons residing in the surrounding 1 mm^3^ volume. By applying these constraints, we maintain the structural rigor of a proofread subcircuit while exploiting the extensive statistical power of the broader cortical network.

**Figure 1.**
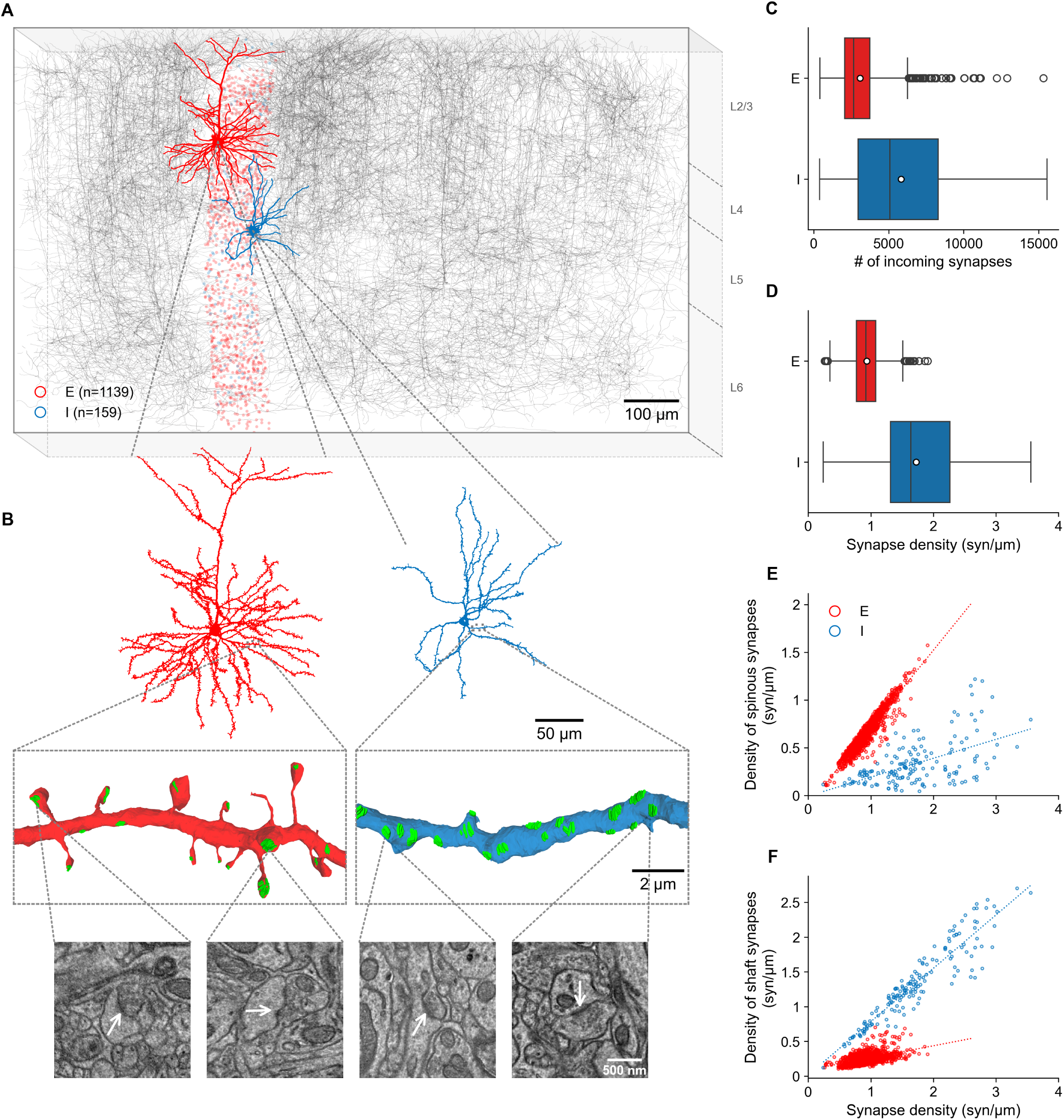
Distinct synaptic connectivity of excitatory and inhibitory neurons. Excitatory neurons receive mostly spine synapses whereas inhibitory neurons receive more synapses on dendritic shafts. **A.** Gray network visualization of the ∼1 mm^3^ mouse V1 circuit, derived from the MICrONS dataset dense reconstruction; 400 neurons were randomly sampled for this display. The proofread micro-column (colored) contains 1,298 neurons (1,139 excitatory, red; 159 inhibitory, blue) after basic structural filtering (see Methods) and is connected by ∼146K intrinsic synapses, receiving a total of ∼4.6M incoming synapses from the whole volume, and making ∼1M outgoing synapses. **B.** *Top:* Zoom-in of two representative neurons from the micro-column shown in A: a layer 2/3 pyramidal neuron (red) and a L5 basket interneuron (blue); only dendrites are shown. *Middle:* Higher magnification views of representative dendritic branches from the neurons shown above. *Bottom:* Representative EM sections illustrating synapses contacting the dendrites shown above; arrows pointing at the post-synaptic density. **C–D.** Quantification of synaptic inputs for excitatory and inhibitory neurons within the whole volume; **C** shows the number of incoming synapses per neuron whereas **D** displays synapse density (synapses/µm). In both panels, the box edges represent the 25th and 75th percentiles, the vertical line denotes the median, and the white circle indicates the mean. Whiskers extend to 1.5× the interquartile range. Mean ± SD values: **C**, excitatory: 3,097 ± 1,714 vs. inhibitory: 5,828 ± 3,507; **D**, excitatory: 0.97 ± 0.25 vs. inhibitory: 1.81 ± 0.68 synapses/µm. **E.** Density of spinous synapses as a function of total synapse density for excitatory and inhibitory neurons. Dotted lines represent linear regressions (excitatory: slope = 0.81, *R*^2^ = 0.9, *p <* 0.001; inhibitory: slope = 0.2, *R*^2^ = 0.3, *p <* 0.001). **F.** Density of shaft synapses as a function of total synapse density. Dotted lines represent linear regressions (excitatory: slope = 0.18, *R*^2^ = 0.28, *p <* 0.001; inhibitory: slope = 0.77, *R*^2^ = 0.87, *p <* 0.001).

### Differences in connectivity among excitatory and inhibitory neurons

Figure 1 illustrates the distinct synaptic connectivity of excitatory and inhibitory neurons in a densely reconstructed cortical circuit from mouse V1 (Fig. 1A). The analysis is based on a proof-read micro-column (Fig. 1A, colored) derived from the ∼1 mm^3^ MICrONS dataset of mouse V1 (Fig. 1A, grey box). The micro-column initially contained 1,352 manually identified neurons, which were filtered to a final dataset of 1,298 neurons with fully resolved structures (1,139 excitatory and 159 inhibitory; see Methods). These neurons are connected through ∼146,000 intrinsic synapses (where both the pre and the postsynaptic neurons are in the sub-circuit), receiving a total of ∼4.6 million incoming synapses from the surrounding tissue (from the 1 *mm*^3^ cut volume and, notably, also from areas outside of it, such as the thalamus) and forming ∼1 million outgoing synapses onto the whole volume (Fig. 1A). Representative reconstructions highlight the striking morphological differences between the excitatory and inhibitory classes (Fig. 1B). Excitatory pyramidal neurons exhibit dendrites densely decorated with dendritic spines, whereas inhibitory interneurons display comparatively smooth dendrites with far fewer spines (Fig. 1B). Zoomed views of representative dendritic branches and corresponding EM sections illustrate that excitatory dendrites receive mostly spine synapses, whereas inhibitory dendrites are contacted primarily by shaft synapses.

Quantitative analysis of synapse number and density shows that inhibitory neurons receive more and denser synaptic inputs than excitatory neurons. On average, inhibitory cells receive almost twice as many incoming synapses, 5,828 ± 3,507 vs. 3,079 ± 1,714, respectively (Fig. 1C) and exhibit roughly two-fold higher synapse density along their dendrites, 1.81 ± 0.68 synapses/µm vs. 0.97 ± 0.25 synapse/µm, respectively (Fig. 1D).

Previous ultrastructural and functional mapping studies have suggested distinct subcellular targeting preferences for excitatory and inhibitory neurons (Kwon et al., 2019), a bias that becomes strikingly evident at scale in the MICrONS dataset. In excitatory neurons, the majority (70%) of synaptic contacts occur on dendritic spines whereas the dendritic shaft is only sparsely decorated with synapses; the variability of synapse density is almost completely explained by variability of spine synapses (Fig. 1E), indicating that connectivity expansion onto excitatory neurons occurs primarily through dendritic spines. This means that, in excitatory neurons with low spine density, the lack of spine synapses is not compensated by an increase in shaft synapses, even though there is more space on the shaft. Conversely, inhibitory neurons receive most (80%) of their synapses directly on the dendritic shaft; their synapse count scales almost entirely through shaft synapses (Fig. 1F), which increases steeply with synapse density. Together, these results strengthen and scale up earlier observations of compartment-specific spine versus shaft connectivity in excitatory and inhibitory cortical neurons (Chen et al., 2012).

These findings point to a fundamental principle of cortical circuits: excitatory neurons integrate inputs through spine-based compartments that may enable more selective and localized dendritic computations, whereas inhibitory neurons receive denser, likely less selective, shaft-based inputs that may perform dendritic computations more globally (see below and Discussion).

### Dendritic spines density correlates with input number and diversity

Figure 2 shows that dendritic spine density is correlated with the richness and diversity of synaptic inputs to the respective neuron. Figure 2A illustrates two reconstructed pyramidal neurons showing the spatial distribution of their presynaptic partners originated within the local micro-column depicted in Figure 1. Presynaptic neurons are distributed throughout the surrounding network and include both excitatory (triangle) and inhibitory (circle) partners. Representative dendritic segments from excitatory and inhibitory neurons further illustrate the structural contrast between spiny and aspiny dendrites (Fig. 2B).

**Figure 2.**
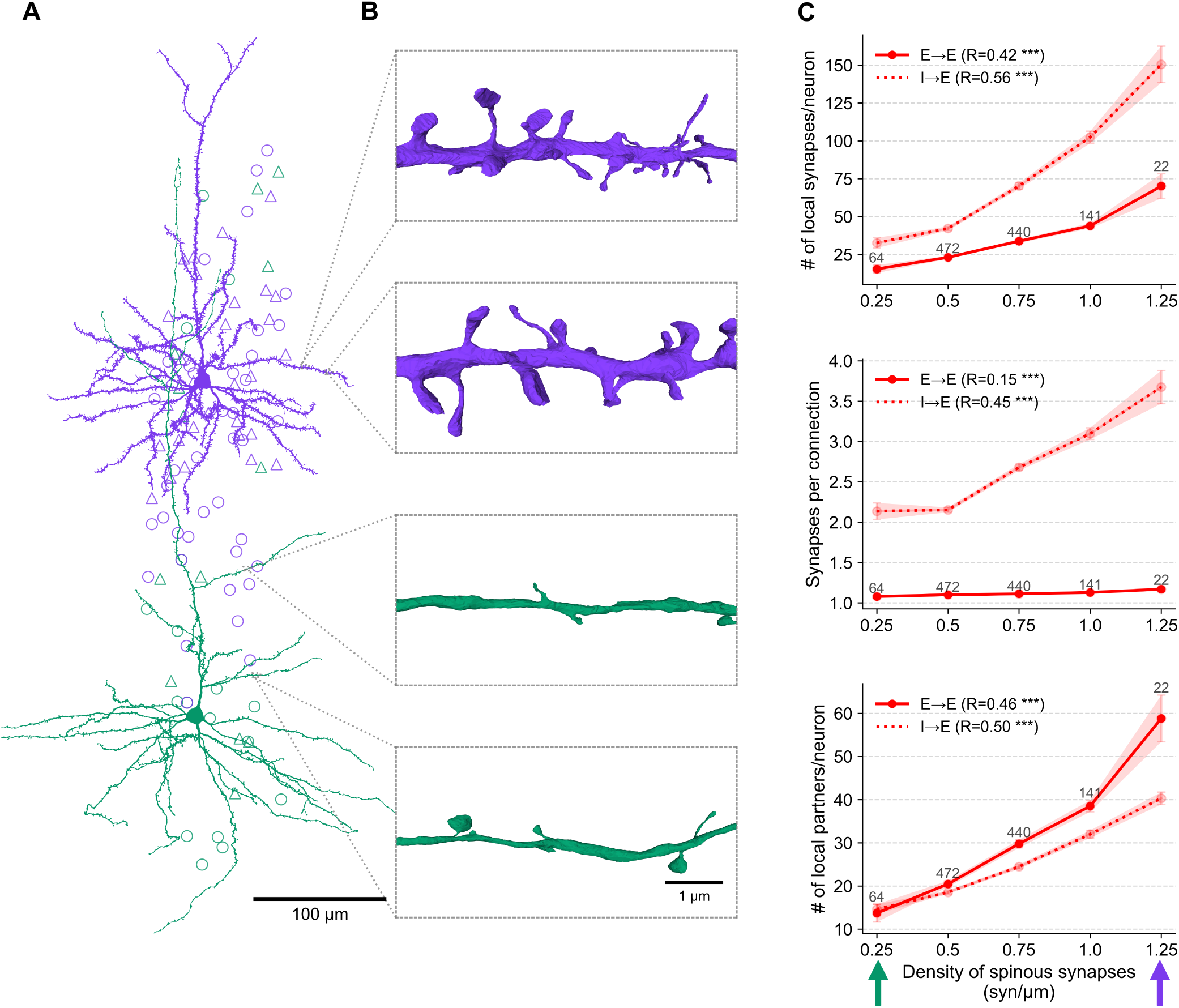
Spine density correlates with local synaptic input richness and diversity. **A.** Example reconstruction of layer 4 (purple) and layer 6 (green) cortical pyramidal neurons with two different spine densities, depicting the spatial distribution of their presynaptic partners’ soma location within the micro-column shown in Figure 1A; triangles for excitatory and circles for inhibitory; only dendrites are shown. **B.** Representative dendritic segments from neurons shown in A, illustrating the contrast between densely and sparsely spiny dendrites of excitatory neurons of different types. **C.** *Top:* Relationship between number of local synaptic inputs per neuron, originating within the micro-column, and total spine density for excitatory neurons. *Middle:* As in top but for the number of synapses per connection (i.e., per presynaptic axon). *Bottom:* As in top but for number of distinct presynaptic partners per neuron. Solid lines represent excitatory-to-excitatory connections and dashed lines inhibitory-to-excitatory connections. Arrows at the bottom illustrate the spine density of the neurons in panel A.

**Figure 3.**
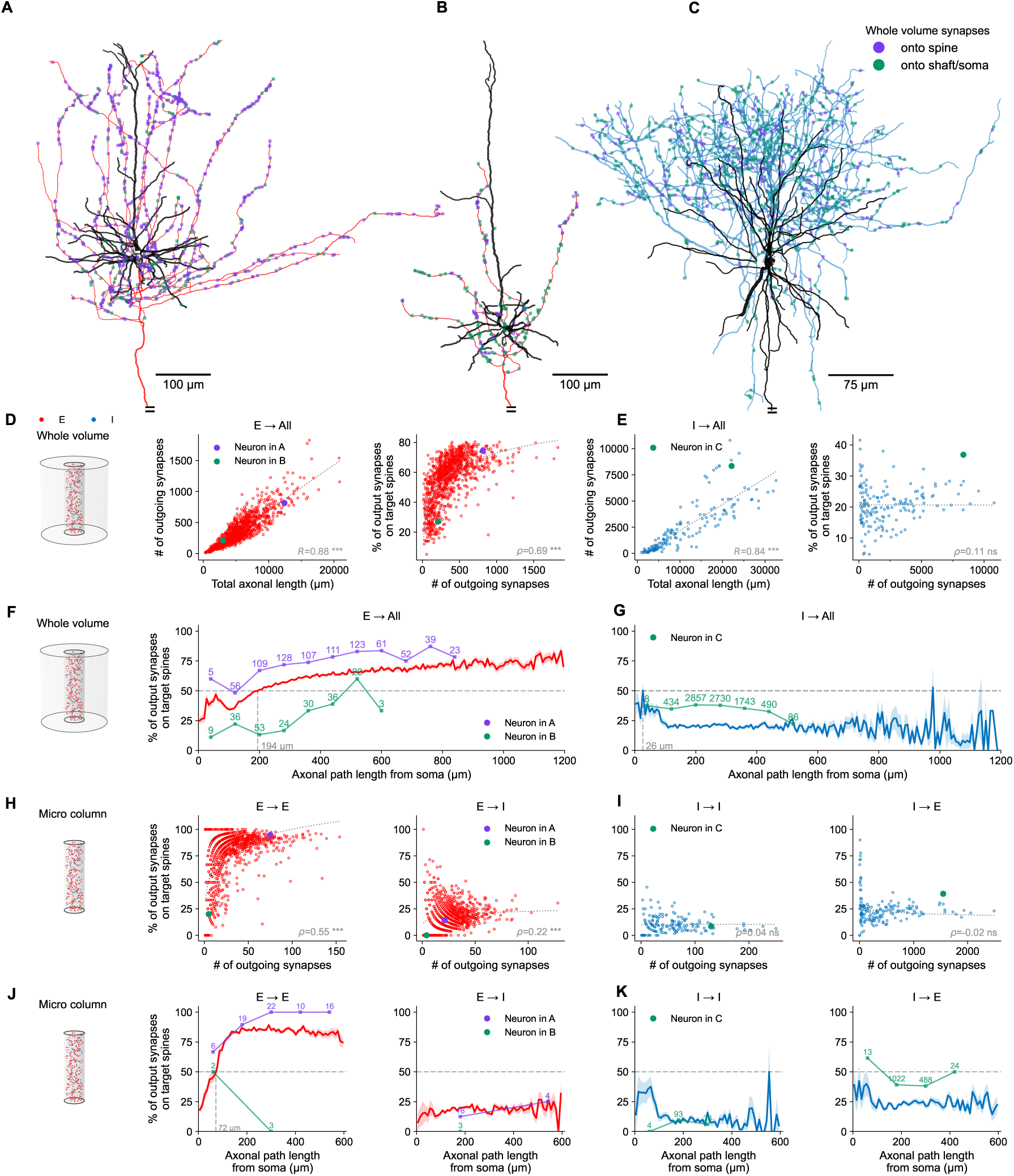
Spine targeting increases with axonal distance from soma in excitatory but not inhibitory neurons. **A.** Reconstructed L5-IT pyramidal neuron with an extensive axonal arbor and a large number of outgoing synapses. Dendrites are shown in black and axons in red, synapses onto spines in purple and shaft/soma synapses in green. **B.** Reconstructed L6-CT pyramidal neuron with a shorter axon and fewer outgoing synapses. **C.** Reconstructed L4 basket interneuron. Dendrites are shown in black and axons in blue. For visualization clarity, 10% of outgoing synapses are randomly sampled and colored as in A–B. **D.** *Left:* Number of excitatory axonal synapses versus total axonal length; dotted line indicates an average density of 7 synapses per 100 µm of axon (*R* = 0.88*, p <* 0.001). Neurons in A and B are highlighted in purple and green, respectively. *Right:* % of excitatory synapses onto target spines as a function of the total number of outgoing synapses (*ρ* = 0.69*, p <* 0.001). **E.** As in D for inhibitory axons. *Left:* Number of synapses versus axonal length (*R* = 0.84*, p <* 0.001), with an average density of 25 synapses per 100 µm. The example neuron in C is highlighted (green). *Right:* % of synapses onto target spines showing no significant dependence of spine targeting on the number of outgoing synapses (*ρ* = 0.11*, p* = 0.15). **F.** Spine targeting as a function of path distance along excitatory axons. Population average (red) and individual neurons from A (purple) and B (green) are shown, with the number of synapses per bin indicated above respective curves. The horizontal dashed line marks the distance at which 50% of synapses are made onto spines. **G.** Same as in F for inhibitory axons. Spine targeting remains low and largely independent of path distance along the axon. **H.** *Left:* Excitatory-to-excitatory (E→E) connectivity within the micro-column shows a strong increase in spine targeting with the number of outgoing synapses (*ρ* = 0.55*, p <* 0.001). *Right:* Excitatory-to-inhibitory (E→I) connectivity shows a weaker relationship (*ρ* = 0.22*, p <* 0.001). **I.** Same as in H for inhibitory axons. Inhibitory-to-inhibitory (I→I) and inhibitory-to-excitatory (I→E) connections show no relationship between the number of outgoing synapses and spine targeting (*ρ* ≈ 0, ns). **J.** Same as in F for the micro-column. *Left:* E→E shows similar increase as in F. *Right:* E→I connectivity shows a lower baseline of spine targeting that remains shallow across axonal path lengths. **K.** Same as in G for the micro-column. *Left:* I→I and *Right:* I→E connections both show that spine targeting remains low and independent of axonal path distance from the soma. Statistical significance for linear correlations was determined using Pearson’s *R* or Spearman’s rank correlation (*ρ*) as indicated.

Across the population of pyramidal neurons, higher spine density is associated with increased local synaptic input originating from within the micro-column (Fig. 2C, top). The number of synapses increases by approximately a factor of 4.5, from 15.4 to 70.1, as spine density increases from 0.25 to 1.25 spines/µm (Fig. 2C). For excitatory inputs this is expected from the overall increase in synapse count associated with increased spine density (Fig. 1E).

Remarkably, we observed a similar increase in number of inhibitory inputs with increase in spine density, although inhibitory synapses largely target shafts (∼80%, see Fig. S5), which means that their abundance could have been independent of spines (Fig. 2C, top, dotted line). For inhibitory presynaptic neurons, the number of synapses that they make per postsynaptic excitatory neuron increases by a factor of 4.6, from 32.6 to 150.5, as spine density increases from 0.25 to 1.25 spines/µm (see Methods).

Interestingly, an increase in spine density is also associated with a larger number of synapses made by individual inhibitory axons onto the postsynaptic cell (i.e., multiple connections). The average number of inhibitory synapses per axon per postsynaptic neuron increases from 2.1 synapses/axon to 3.6 synapses/axon with an increase in spine density of the postsynaptic excitatory neuron (Fig. 2C, middle, dotted line). In contrast, excitatory axons typically form only a single synapse per axon over the target neuron, largely independent of spine density (Fig. 2C, middle, filled line).

The lower redundancy of excitatory compared to inhibitory inputs could be the statistical result of the lower count of local excitatory synapses, combined with the smaller population of potential presynaptic inhibitory partners. However, this means that the increase in excitatory synapse density resulting from higher spine density is directly translated into an increase in the number of distinct presynaptic partners, which rises from 13.7 partners to 58.8 excitatory partners as spine density increases from 0.25 to 1.25 spines/µm (Fig. 2C, bottom). Thus, in excitatory neurons, dendrites with higher spine density receive synaptic input from a larger and more diverse set of presynaptic neurons.

Equivalent analyses across the population of inhibitory neurons reveal a different organizational principle (Fig. S1C; representative morphologies depicted in Fig. S1A–B). Although inhibitory neurons receive substantially more local synaptic input overall (the majority of neurons receive between 100 and 200 excitatory synapses across the observed spine-density range, with only 17 neurons receiving between 200 and 400 synapses). For inhibitory neurons, the number of inhibitory synapses/neuron remains roughly constant (56.9 synapses/neuron) and largely independent of spine density (Fig. S1C top, dotted line). Moreover, the scaling of multiple synapses from individual presynaptic axons diverges sharply between cell types. While excitatory neurons show a pronounced increase in multi-synaptic connections from inhibitory inputs, inhibitory neurons exhibit the inverse pattern: increased spine density is instead associated with a slight increase in multiple synapses formed by individual excitatory axons (E→I, peaking around 1.5 to 2 synapses per connection, Fig. S1C middle), whereas connections from individual inhibitory axons (I→I) show little to no increase. Furthermore, higher spine density in these neurons is specifically associated with a larger number of excitatory presynaptic partners (increasing from an average of ∼85 to ∼139), while the number of inhibitory presynaptic partners remains approximately constant (21 partners per neuron, Fig. S1C, bottom). This relationship holds true even when normalizing for total dendrite length (Fig. S2), indicating that these overarching trends are not merely a byproduct of larger dendritic trees but represent a specific connectivity logic.

These findings refine our understanding of how dendritic structures mediate connectivity. In excitatory neurons, spines clearly serve to expand network sampling. In inhibitory neurons, the architecture is different: shaft synapses are the main driver of input diversity (Fig. 1); yet spines still maintain a measurable impact on connectivity by facilitating more (excitatory) partners.

Comparison of presynaptic partner counts for excitatory and inhibitory neurons (Fig. 2C–D, bottom) together with the results of Figure 1 reveal a connectivity principle in local cortical circuits. In both cell classes, dendritic spine density acts as a structural predictor for the richness and diversity of synaptic input, providing quantitative validation for the connectivity and diversity hypothesis (Yuste, 2011). In excitatory neurons, this relationship is particularly potent, as spine density strongly predicts both the number and diversity of local presynaptic partners. While inhibitory neurons receive denser shaft-based inputs and more presynaptic partners overall, they still follow this “spine-for-diversity” logic specifically for their excitatory inputs. Although this dependence on spine density corresponds to a roughly 4-fold increase in connections for excitatory neurons, the consistent trend across both populations suggests that spines are the primary morphological tool used by cortical neurons to broaden their sampling of the surrounding network.

### Length of excitatory axons correlates with synaptic output number and spine targeting

After describing and characterizing a large variability of spine vs. shaft preference with respect to origin of the input synapses, we next considered neuronal outputs. Specifically, we examined the relationship between axonal extent and synaptic output in reconstructed cortical axons. Previous topological analyses of neocortical microcircuitry have demonstrated that neurons with extensive, highly branched axons effectively function as dense “outgoing hubs” that broadly distribute signals across the network (Gal et al., 2017). Our volumetric data analysis provides direct, large-scale structural evidence for this architecture. Representative examples of reconstructed axons are shown in Figure 3A–C; a long-range L5-IT pyramidal neuron forming large numbers of synapses (Fig. 3A); L6-CT pyramidal neuron with a shorter axon and with much less output synapses (Fig. 3B); and an axon of a basket cell forming dense output synapses (Fig. 3C).

Analysis of the connectivity of excitatory axons, originating within the micro-column but projecting both in and outside of the micro-column, shows that the total number of outgoing synapses correlates linearly with axon length (Fig. 3D, left; Pearson *R* = 0.88), with a slope of 7 synapses/100 µm (one synapse per 14 µm) for excitatory axons, in agreement with (Loomba et al., 2022; Piazza et al., 2025). Interestingly, and in agreement with Loomba et al. (2022) and Schmidt et al. (2017), the ultrastructural target of these output synapses changes systematically with increasing synaptic output number; the fraction of synapses onto dendritic spines increases with the number of outgoing synapses (Fig. 3D, right; Spearman *ρ* = 0.69). Namely, excitatory neurons with longer axons not only make more output synapses in total, but also exhibit a stronger preference for spine targeting. Using the micro-column where the identification of both pre- and post-synaptic neurons is known, we next examined whether the relationship between synaptic output and spine targeting depends on postsynaptic cell (E or I) type. Excitatory-to-excitatory (E→E) connections show a strong increase in spine targeting with the number of outgoing synapses (Fig. 3H, left; *ρ* = 0.55). In contrast, excitatory-to-inhibitory (E→I) connections show a weaker dependence (Fig. 3H, right; *ρ* = 0.22).

In this context, it should be noted that we use the terms “spine targeting” or “axonal preference for spines” as a descriptive shorthand for our observations; we do not imply that the mechanisms underlying the observations lie exclusively on the presynaptic side, nor do we discount the active roles of the spines in the formation of a synapse. To determine whether distance-dependent spine preference reflects spatial organization along the axon, we quantified spine targeting as a function of axonal path distance from the soma. Across the excitatory axons, the fraction of synapses onto spines transitions from a low baseline near the soma to a spine-dominated regime at larger distances (Fig. 3F, red curve). This progression exhibits a transient dip around 100 µm; analysis of the underlying synapse distributions reveals that this dip is driven by a dense, localized zone of shaft-directed synapses onto nearby inhibitory interneurons (Fig. S4B).

Notably, spine targeting transitions from a mixed (shaft/spine) regime near the soma to a spine-dominated regime at larger distances (Loomba et al., 2022; Schmidt et al., 2017). We found that, at the population level, the distance at which spine targeting reaches 50% is ∼200 µm (Fig. 3F, dashed line, see Methods), which provides a characteristic length scale for this transition. However, individual neurons exhibited substantial diversity. A large subset of neurons preferred spines all along their reconstructed axon, remaining continuously above this 50% threshold (37%), while a very small fraction never reached this threshold (0.1%). For the remaining neurons that did have this transition distance, the crossing occurred at a median distance of 83 µm (IQR: 49.1–114.5 µm), with distinct distributions across different morphological cell types (Fig. S7, see Methods).

Given that local inhibitory targets primarily receive synapses on their dendritic shafts while distal excitatory targets integrate inputs via spines, this spatial transition is naturally related to the distance-dependent shift in target cell-type preference described in these prior studies (Schmidt et al., 2017; Bodor et al., 2025). However, we demonstrate that a shift in postsynaptic cell type alone cannot fully account for this phenomenon. Crucially, when the local inhibitory population is removed and the analysis is strictly confined to the excitatory-to-excitatory (E→E) subnetwork, the localized dip disappears, and the progressive increase in spine targeting remains highly pronounced (Fig. 3J, left). In contrast, for excitatory-to-inhibitory (E→I) connections, as well as all inhibitory outputs (I→I and I→E), spine targeting stays at a lower baseline and remains largely independent of axonal path distance from the soma (Fig. 3J, right; Fig. 3K). This indicates that the progressive preference for dendritic spines is a subcellular organizing principle of excitatory axons, rather than a mere by-product of cell-type targeting. In fact, the result might indicate the reverse: that the shift in target cell-type preference is a side effect of (or implemented as) a shift in spine preference. In that case, cells with an overall lower preference for spines would also exhibit a lower preference for excitatory targets. We found that this is indeed the case. When examining the local connectivity within the micro-column, an excitatory neuron’s preference for targeting other excitatory partners (quantified as the ratio of E to E+I partners) is only weakly correlated with its total number of outgoing synapses (Spearman *ρ* = 0.20). However, this preference is robustly predicted by the neuron’s overall preference for targeting dendritic spines (Spearman *ρ* = 0.56; Fig. S6).

The observed increase in the probability, *p*(*x*), of spine targeting with axonal path length from soma can be captured by a simple model:

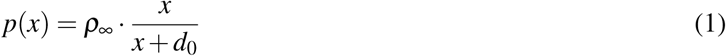

where *x* is the axon’s path distance from soma to the synapse (measured in µm), *ρ*_∞_ is the asymptotic spine-targeting probability, and *d*_0_ is a characteristic length scale governing the transition from the initial regime which is biased towards the dendritic shaft to the asymptotic regime, dominated by spines.

We tested the fit of this equation to our data; the results are shown in Figure S4. This showed two basic results: first, the fit is robust against the number of spatial bins used; second, the model works best at the population level. The natural variation and random branching within a single axon introduce greater noise and make the fit much less reliable compared to an entire population. We found that the asymptotic spine-targeting probability (*ρ*_∞_) remains consistently high (ranging from 0.82 to 0.93), but the characteristic length scale (*d*_0_) varies drastically depending on the target population. For these presynaptic excitatory neurons, the characteristic scale, *d*_0_ is 124 µm when considering all available targets across the entire volume (E→All) and 148 µm for all target neurons (E + I) within the micro-column (E→E&I). This scale decreases to 28 µm when considering strictly excitatory targets within the micro-column (E→E) (Fig. S4). This indicates that the transition to a spine-targeting preference happens over a much shorter distance when axons are restricted to an excitatory-only network.

On the level of individual neurons, we found substantial variability in the overall fraction of spine synapses targeted by their axon (Fig. 3F, purple vs. green), but the increase of the probability to target a spine with path distances from soma still holds (Eq. 1). This indicates that the population-level increase is not the consequence of averaging over axons of different lengths. Neither is the increase in spine preference for longer axons explained by longer axons simply having more mass at longer distances (Fig. S8). To determine if distance alone could account for this trend, we applied our distance-dependent simple model (fitted to the population average, Fig. S4) to the empirical synapse distances of each individual neuron. By calculating a predicted global spine preference based purely on the spatial distribution of each neuron’s synapses, we found that distance-based rules alone cannot replicate the wide cell-to-cell variance or the full magnitude of the increase in spine preference seen in neurons with many outgoing synapses. In other words: higher spine preference for longer axons and higher spine preference for distal axon segments are two separate, concurrent phenomena.

### Length of inhibitory axons correlates with synaptic output number but not spine targeting

In contrast to excitatory axons, inhibitory axons exhibit a different organization principle. Although axonal length strongly correlates with the number of outgoing synapses, with an average of 25 synapses/100 µm (one synapse/4 µm, Fig. 3E, left; Pearson *R* = 0.84), the fraction of synapses onto spines does not depend on the number of axonal synapses (Fig. 3E, right; Spearman *ρ* = 0.11). Consistent with this, spine targeting of inhibitory axons remains low and largely independent of path distance along the axon (Fig. 3G), indicating the absence of a distance-dependent transition from shaft-to-spine preference. Thus, inhibitory axons do not exhibit the progressive shift toward spine targeting observed in excitatory axons.

When analyzing the micro-column subset, in inhibitory neurons, no significant relationship is observed for either inhibitory-to-inhibitory (I→I) or inhibitory-to-excitatory (I→E) connections (Fig. 3I), reinforcing the distinct targeting rules governing inhibitory connectivity. Further unpacking these divergent strategies, we find that the capacity to target spines directly drives the expansion of outgoing connectivity for excitatory neurons (Fig. S5). Their axons appear to establish a baseline supply of shaft-targeting synapses, with variations in overall output density determined almost completely by the addition of spine-targeting synapses. Inhibitory neurons operate under a fundamentally different logic; while their overall density of outgoing synapses is highly variable, they maintain a relatively constant spine-to-shaft ratio of approximately 20% (Fig. S5), scaling their output almost exclusively through shaft-targeting synapses.

To ensure that the observed relationship between axonal extent and spine targeting is not an artifact of incomplete axonal reconstruction, we performed a control analysis restricted to a subset of the most rigorously proofread axons, both inside and outside the micro-column (Fig. S3). A potential confound in large-scale connectomic datasets is that partially reconstructed axons might artificially appear both short and uniquely biased in their synaptic targeting, thereby driving a spurious population-level trend. To rule out this possibility, we limited our analysis strictly to a highly curated subset of axons with the highest level of manual proofreading (referred by the Allen Institute as “axon_fully_extended”, see Methods). This subset consists of 272 excitatory and 7 inhibitory neurons from within the micro-column, alongside an additional 42 excitatory and 126 inhibitory fully proofread neurons located outside the micro-column. These fundamental trends remain robust within this combined group. Total synaptic output continues to correlate linearly with total axonal length, and the fraction of spine-targeted synapses for excitatory axons still systematically increases with both total synaptic output (Fig. S3A) and path distance from the soma (Fig. S3C). This suggests that the progressive transition toward dendritic spines reflects a genuine biological feature of extended excitatory axons, rather than a mere byproduct of truncation or reconstruction artifacts; this is further supported by the results in different cortical systems reported in Loomba et al. (2022) and Schmidt et al. (2017).

Together, these results reveal principles of cortical excitatory connectivity: excitatory axons with a greater total number of outgoing synapses exhibit a higher overall tendency to target dendritic spines, and this spine targeting also increases in a distance-dependent, saturating manner as the axon extends from the soma. Because distance alone cannot fully account for the higher spine targeting observed in neurons with extensive synaptic outputs, these likely represent concurrent but distinct structural phenomena. The spatial transition from early shaft-biased targeting to a spine-dominated regime is well captured by a simple model (Eq. 1) defined by a single transition scale *d*_0_, reflecting the extent of the excitatory axon required to overcome early synaptic targeting bias towards the dendritic shaft. In contrast, inhibitory neurons do not exhibit such a transition, indicating fundamentally different structural rules for excitatory and inhibitory axonal targeting.

### Shared-input innervation is correlated with spine targeting

The organization of cortical circuits relies heavily on higher-order connectivity motifs to shape network function. A key principle in this domain is the ‘common neighbor’ rule (Perin et al., 2011), which establishes that the probability of a synaptic connection between cortical neurons is strongly predicted by the number of connections they share. While this collaborative architecture was originally identified functionally in cortical slices (Perin et al., 2011; Song et al., 2005), its underlying ultrastructural logic, i.e., its relation to spine targeting and spine anatomy has remained unexplored. Therefore, we next investigated the structural rules governing shared presynaptic input across the whole dataset.

Towards this end, we define “shared input strength” - a metric designed to measure the collaborative nature of each postsynaptic neuron, based exclusively on its incoming connectivity. We define the shared input strength of a neuron *N* as the mean outdegree of the presynaptic population of *N*. The measure is directly related to the concept of common neighbors as follows: Consider a neuron *P* that is presynaptic to *N*. Each neuron innervated by *P* is a neuron that shares *P* as a common (input) neighbor with *N*. Therefore, a high outdegree of *P* translates directly to an increased mean number of common neighbors for pairs of neurons including *N* (see also Appendix). For example, both the red and blue neurons in Fig. 4A are innervated by three presynaptic neighbors (gray with black outline), but for the red neuron these neighbors have a higher outdegree. Consequently, the red neuron shares all three of them as common neighbors with each of various peer neurons in the vicinity (gray circles), while the blue neuron only shares one of them with each.

**Figure 4.**
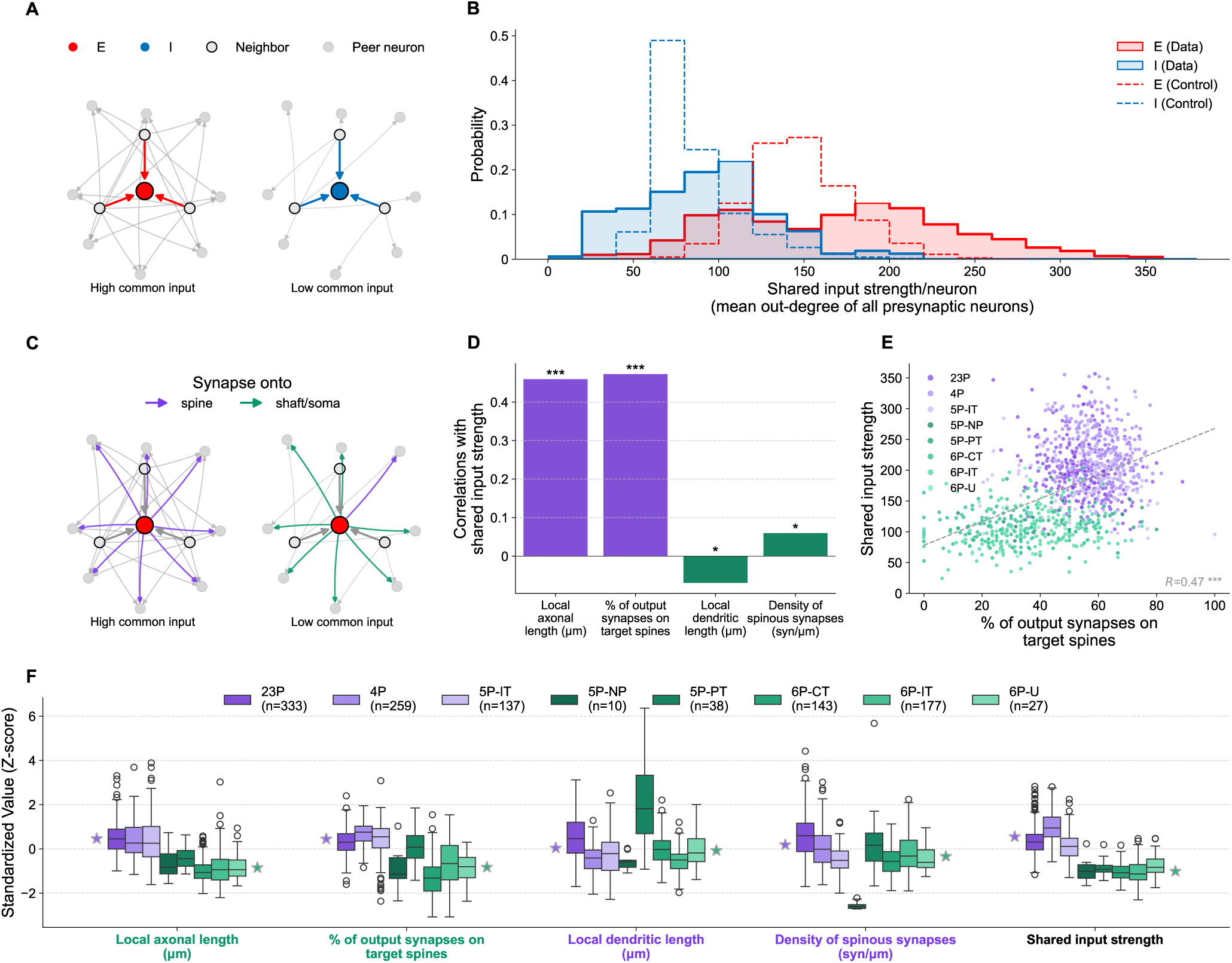
Shared-input innervation correlates with spine targeting. **A.** Schematic illustration of the sharing property: excitatory (E, red) and inhibitory (I, blue) neurons each receive input from 3 common presynaptic neighbours (gray circles with black outline) that are evaluated against surrounding peer neurons (solid gray circles). In the case of the E-neuron, its 3 presynaptic partners are high-out-degree nodes projecting broadly across the network, whereas the presynaptic partners of the I-neuron do not. Consequently, the E-neuron has a high tendency to share input with the rest of the network (shared input strength: 9). This is lower for the I-neuron (shared input strength: 3.6) **B.** Distribution of the shared input strength/neuron in the whole population. The excitatory (solid red) and inhibitory (solid blue) populations are compared against null distributions from 100 degree-preserving “blocked” configuration (CFG) models (dashed lines). Excitatory and inhibitory networks exhibit significantly distinct sharing properties (Mann-Whitney *U* = 152,851, *p <* 0.001; Kolmogorov-Smirnov *D* = 0.55, *p <* 0.001). The shared input strength for excitatory neurons is significantly more broadly distributed compared to the respective null distribution (Kolmogorov-Smirnov *D* = 0.34, *p <* 0.001; Mann-Whitney *U* = 79,810,238, *p <* 0.001), whereas the shared input strength of inhibitory neurons tends to be lower than the respective null model (Kolmogorov-Smirnov *D* = 0.25, *p <* 0.001; Mann-Whitney *U* = 1,439,959, *p* = 0.0025). **C.** Schematic illustrating the relationship between the shared input strength and spine targeting. *Left:* a highly sharing neuron preferentially targets dendritic spines (purple arrows). *Right:* low sharing neuron tends to target the dendritic shaft (green arrows). **D.** Correlation between the shared input strength against four features: the two left panels in purple display output features, while the two right panels in green display input features. **E.** Correlation between the neuron’s shared input strength and its spine preference (Pearson *R* = 0.47, *p <* 0.001). **F.** Z-scores of the features shown in E across different excitatory cell types. 2/3P, L4P, and 5P-IT populations (in purple) exhibit the highest shared input strength and spine targeting preferences; deep-layer neurons and neurons providing cortical output (in green) display lower shared input strength and less spine-targeting preference. The purple and green stars indicate the mean per feature for all cell types in their respective colors.

Shared input strength is a measure of how much the input that *N* receives is also received by other neurons. We interpret such sharing of inputs as a propensity of *N* to collaborate with other neurons in the representation of information. This is because it describes a setup where the same, or strongly overlapping information is received by multiple neurons that may then represent the information collaboratively in the form of a population code.

We investigated the properties of shared input strength in the source data and then its relation to neuronal spine anatomy. First, we found that input sharing is overall more substantial among E-neurons (Fig. 4B, red) than I-neurons (Fig. 4B, blue, Mann-Whitney *U* = 152,851, *p <* 0.001; Kolmogorov-Smirnov *D* = 0.55, *p <* 0.001). However, the most prominent characteristic of the E-population is the structure of its variability (red line in Fig. 4B) compared to a degree-preserving configuration (CFG) control model (Fig. 4B, dashed red line), E-neurons exhibit a significantly wider distribution of shared input strengths than expected by chance (Kolmogorov-Smirnov *D* = 0.34, *p <* 0.001; Mann-Whitney *U* = 79,810,238, *p <* 0.001). We would like to stress that a degree-preserving control is very strong for a degree-based measure, such as shared input strength. Additionally, to rule out differences in average outdegree between E and I populations as a confounding factor, this CFG model shuffles each block of the matrix (i.e., EE, EI, IE, II) separately. This comparison reveals that, while I-neurons tend to share fewer than predicted by the respective null model (Fig. 4B, dashed blue line, Kolmogorov-Smirnov *D* = 0.25, *p <* 0.001; Mann-Whitney *U* = 1,439,959, *p* = 0.0025), the E-population is characterized by a highly heterogeneous sharing architecture. Similar heterogeneity in graph-based measures has been described in this dataset before and speculatively linked to efficient neural codes (Santander et al., 2025). In this case, it is strong indication of selective connectivity; specifically, that the subpopulation of excitatory neurons with unexpectedly high shared input strength is selectively targeted by neurons with a high outdegree.

To further understand the nature of this targeting within the excitatory population, we evaluated how shared input strength relates to other fundamental morphological and synaptic features. To our surprise, we found the strongest correlations of shared input strength for features of a neuron’s outgoing connectivity instead of its incoming connectivity. Specifically (Fig. 4C), “high-common-input” excitatory neurons preferentially direct their outgoing projections (purple) toward dendritic spines, whereas low-sharing neurons tend to target shafts (green). Indeed, we found that a neuron’s tendency to share presynaptic inputs correlates most strongly with its preference for targeting dendritic spines and with its local axonal length (the total axonal length confined only to the micro-column), surpassing the correlations with the density of the neuron’s spinous synapses, or its local dendritic length (Fig. 4D). Thus, simply being a neuron densely covered with spines is insufficient, as such, for it to be a highly sharing neuron. What is unexpected about this robust structural link between the sharing strength and the axonal length and the tendency to target spines is that the former is fundamentally a property of the input (dictated by the connectivity and out-degree of presynaptic partners), whereas the latter are strictly output features. This solidifies the specific structural link between collaborative input integration and spine-directed synaptic output. Furthermore, this indicates that neurons receiving inputs from neurons with long axons - and hence a preference for spines (Fig. 3H) - have long axons and spine targeting preference themselves. Such a tendency for similar neurons to connect to each other is captured by the term *assortativity*. In this case similarity is based on outdegree, resulting in a form of “rich club” (Van Den Heuvel and Sporns, 2011), but also on axonal spine targeting. Specifically, the assortativity score based on the density of spine-targeting synapses on the axon was 0.28, and the one based on outdegree was 0.2 (not shown).

Figure 4E shows the relationship between the shared input strength and spine targeting. This scatter plot reveals a robust correlation (*R* = 0.47) and highlights an apparent structural bimodality within the population. Indeed, the distribution of shared input strength in Figure 4E and in the histogram of Figure 4B both display two distinct subpopulations. This is confirmed statistically by applying Hartigan’s Dip Test (Hartigan and Hartigan, 1985), which indicated strong evidence against unimodality of shared input strength (Dip statistic = 0.017, *p* = 0.025; see Methods). Notably, this structural divide adheres closely to established cell-type identities from the MICrONS dataset: we colored the high-sharing, spine-preferring population in purple (2/3P, 4P, and 5P-IT) and the remaining populations in green (5P-NP, 5P-ET, 6P-CT, 6P-IT, 6P-U). For clarity, we excluded the ‘unsure E’ (*n* = 4) and ‘WM-P’ (*n* = 11) E neurons from Fig. 4E–F.

To further contextualize the axonal strategies driving the spine preference shown in Figure 4E, we directly compared the densities of outgoing spinous versus shaft synapses across these specific subpopulations (Fig. S12). This whole-volume assessment specifically captures the complete set of approximately 1M outgoing synapses, which project both within and outside the local micro-column. Building on the broad output scaling differences between excitatory and inhibitory neurons established earlier (Fig. S5), we found that the bimodality in excitatory spine preference is driven by distinct subcellular targeting strategies. For each neuron, Fig. S12 plots the density of spine-targeted synapses against the density of shaft-targeted synapses along its axon. In excitatory neurons the two densities vary largely independently (Fig. S12, left): the highly collaborative populations (purple in Fig. 4E) not only target spines more than other neurons, but also spread widely along the outgoing spine-density axis while their shaft density remains comparatively constant. In the low-sharing populations (green), the pattern is inverted, with outgoing shaft density the more variable of the two. For inhibitory neurons, by contrast, spine and shaft densities varied together, leading to strong correlations even across neuron classes (Fig. S12, right). The inhibitory result is consistent with an axon that forms synapses at a variable density, but on any subcellular structure it encounters, without preference. The independence of the two densities in excitatory neurons instead indicates that the ability to target or attract spines is an axon-specific property that varies across classes and individual neurons.

To fully characterize these relationships, we expanded our analysis to evaluate the correlations between shared input strength and structural features across the full network (Fig. 4F, Fig. S10). Furthermore, we also performed this analysis solely on the excitatory sub-network (E-E connectivity; Fig. S9). Consistent with our main findings, output features of the neuron consistently exhibited the strongest correlations with sharing strength in both network contexts. Noteworthy is that in the E-E sub-network a neuron’s spine density - an input feature - emerges as a substantially better predictor of its sharing strength than it is in the full network. Still, even in this E-E subnetwork, spine density remains secondary to the predictive power of spine-targeting output. Finally, to verify that the core trends observed are not driven primarily by the highly curated subset of completely reconstructed axons, analyses were repeated after excluding the neurons with the highest level of manual proofreading (consisting of 272 excitatory and 7 inhibitory neurons from within the micro-column, Fig. S11). Our results remained highly consistent within this partially axon-extended population, with output features continuing to serve as the strongest predictors of a neuron’s shared input strength.

By comparing standardized values (Z-scores) for these metrics across all cell types, a clear layer-specific hierarchy emerged (Fig. 4F). In agreement with previous results (Pedigo et al., 2026), our visualization confirms that the purple-labeled populations exhibit the strongest preference for targeting dendritic spines alongside greater local axonal lengths. Our novel connectivity analysis reveals that these same spine-preferring populations also exhibit the highest propensity for sharing presynaptic partners. In contrast, deeper layer neurons demonstrate both the expected reduction in spine targeting (Pedigo et al., 2026) and noticeably lower sharing properties.

This anatomical distribution aligns intuitively with the canonical flow of cortical information processing (Douglas and Martin, 2004): highly collaborative, spine-preferring connectivity is densely enriched in the primary input (Layer 4) and local processing (Layer 2/3 and L5-IT) layers, whereas layers traditionally associated with broader cortical output (Layer 6) participate less heavily in this specific, local sharing motif.

In summary, an unexpected outcome of this topological analysis is the apparent link between the topology of a neuron’s inputs in the graph representation of a connectome and the subcellular targeting of its outputs. Variability on both sides of this correlation was larger than expected by chance, indicating presence of a mechanism that goes beyond mere stochasticity. The differences aligned largely with differences between excitatory neuron classes, but individual variability within a class was also observed. The observation that a structural *input* feature, a neuron’s tendency to share its input with the network, correlates so closely with its *output* behavior (its preference for innervating dendritic spines) suggests a highly organized “input-to-spine/output-to-spine” organization rule (see Discussion).

### Connectivity patterns are maintained at different circuit spatial scales

To determine whether the observed connectivity patterns are specific to the dimensions of the micro-column or represent fundamental organizational principles of cortical circuitry, we performed a conceptual sub-sampling experiment restricted strictly to excitatory neurons in the micro-column (Fig. 5). By creating nested subnetworks representing fractions of the total volume, ranging from the full micro-column (1,139 E neurons) to progressively smaller scales of 0.5 (771 E neurons), 0.25 (398 E neurons), 0.125 (213 E neurons), and 0.0625 (122 E neurons), we tested the stability of our above conclusions across varying spatial boundaries. We found that the core results of Figures 2–4 remained remarkably consistent across all sub-volumes. We found that the number of local partners, both excitatory (Fig. 5B) and inhibitory (Fig. 5C) consistently increases with increase in spine density of the postsynaptic excitatory neuron. This distinct rising trend is present across all tested subnetwork scales, replicating the primary population-level results of Figure 2C. Similarly, when examining the preference for targeting spines as a function of the number of outgoing synapses (Fig. 5D, re-creating the analysis in Fig. 3H), we observed that a positive correlation is generally maintained across the subnetworks. However, the strength of this correlation noticeably decreases as the subnetwork size shrinks. This attenuation is expected, as the progressive reduction in volume naturally leads to a smaller overall network and a corresponding drop in the number of outgoing synapses available for analysis.

**Figure 5.**
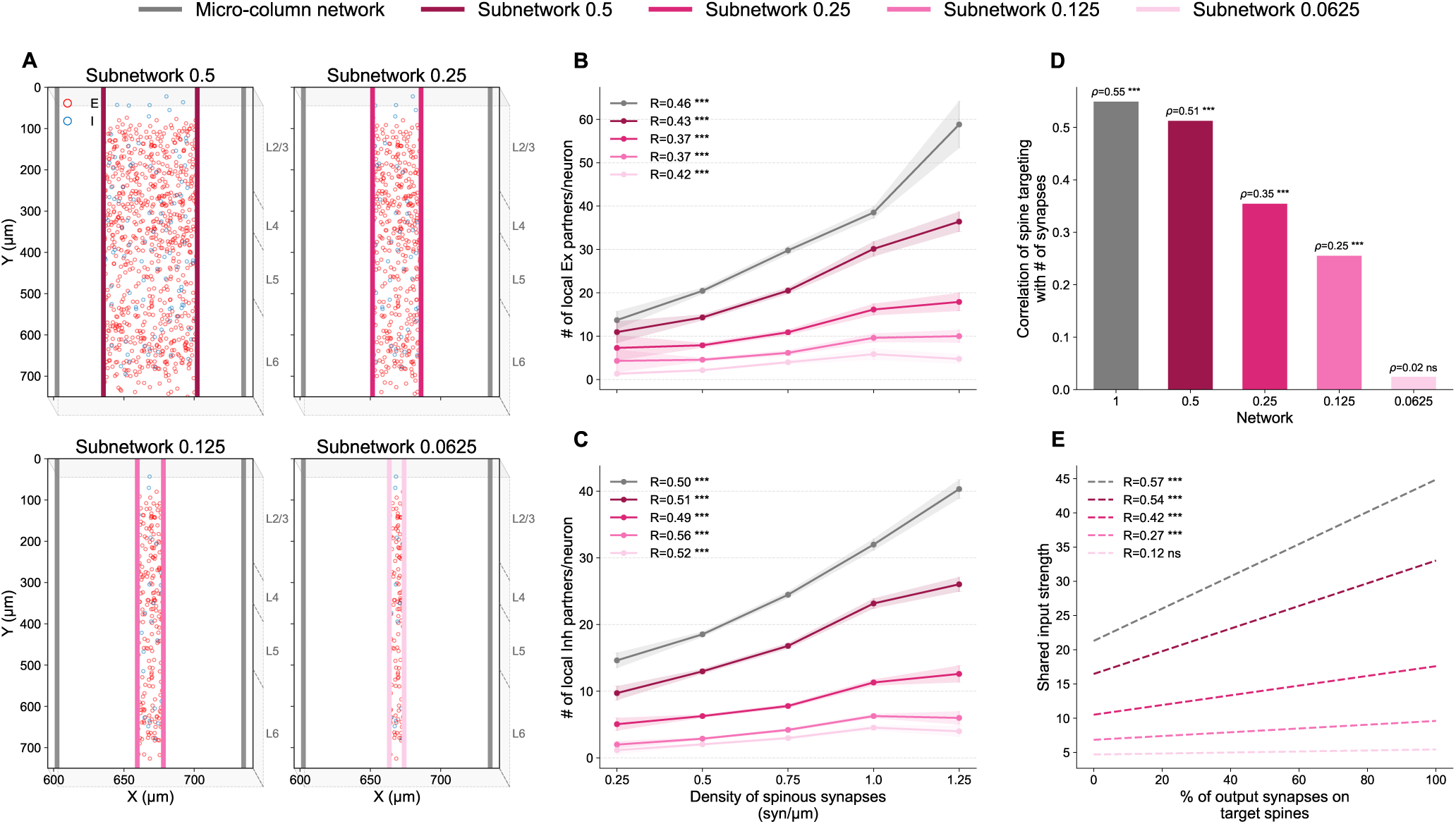
Scale-invariance of connectivity patterns across nested micro-columns. **A.** Illustration of a series of nested subnetworks from the original micro-column, whereby the *X* dimension is scaled by 0.5, 0.25, 0.125, and 0.0625, with *Y* (depth) and *Z* remaining constant. Connectivity and other features within each subnetwork are calculated based solely on neurons and synapses within the respective new boundaries. **B–C.** The number of unique local excitatory (B) and inhibitory (C) partner neurons for postsynaptic excitatory neurons plotted against spine density; same analysis as in Figure 2C. Color-coded lines correspond to the different subnetwork scales; grey for the original column. **D.** Spearman correlation (*ρ*) between the percentage of output synapses on target spines and the number of outgoing synapses for each subnetwork size, corresponding to the analysis in Figure 3D. **E.** Regression lines demonstrating the correlation between a neuron’s shared input strength and its percentage of output synapses on target spines across subnetworks scales, corresponding to the analysis in Figure 4E.

Finally, we examined the relationship between shared input strength within the excitatory population across nested scales (Fig. 5E and Fig. S13). Consistent with our observations in the full network (Fig. 4E and Fig. S9), we found that neurons sharing more presynaptic inputs generally tend to exhibit a greater preference for targeting spines in their output connections. The full underlying data distributions for these subnetwork regressions are available in Figure S13.

Similar to Figure 5D, while the trends remain positive across the larger sub-volumes, the strength of the correlation gradually weakens with decreasing scale. Ultimately, at the smallest subnetwork tested (scaled by 0.0625), the correlations shown in Figure 5D–E lose statistical significance (Spearman *ρ* = 0.02, *p* = n.s. and Pearson *R* = 0.12, *p* = n.s., respectively). We attribute this decay in correlation strength to the sparse sampling within this minimal spatial volume, which is reduced to just 122 excitatory neurons and proportionally fewer synapses, rather than a fundamental break in the connectivity rule.

### Dendritic spines preferentially connect neuronal ensembles

Based on our structural analyses we predict that dendritic spines provide a physical substrate for coordinated network activity: spine density reflects the diversity of a neuron’s inputs (Fig. 2), and neurons that share presynaptic input with the rest of the network preferentially route their own output onto spines (Fig. 4). If this architecture indeed organizes network-level cooperation, its signature might be detectable directly in groups of neurons that are functionally coactive (defined as neuronal ensembles). We tested this prediction using *in vivo* two-photon calcium imaging paired with the same anatomical volume (MICrONS Consortium, 2025; see Methods).

To identify candidate functional ensembles, we applied an ICA-based detection algorithm (Lopes-dos Santos et al., 2013) to trial-residual activity computed by subtracting each neuron’s average response to repeated natural-movie clips, isolating co-fluctuations not explained by shared stimulus tuning (see Methods). Figure 6A shows an example ensemble of four coactive L2/3 pyramidal neurons across six repeated oracle movie clips (MICrONS Consortium, 2025; see Methods) (left). Across the recorded excitatory population with matched structural and functional data (476 neurons, median 41 recorded neurons per scan), this approach yielded 52 candidate ensembles across 13 of 16 recording sessions, ranging in size from 2 to 6 neurons (Fig. 6A, right). Note that this recorded subset under-samples the deep-layer types relative to the superficial ones (Fig. S15A), a limitation of the calcium data that applies to all analyses below.

**Figure 6.**
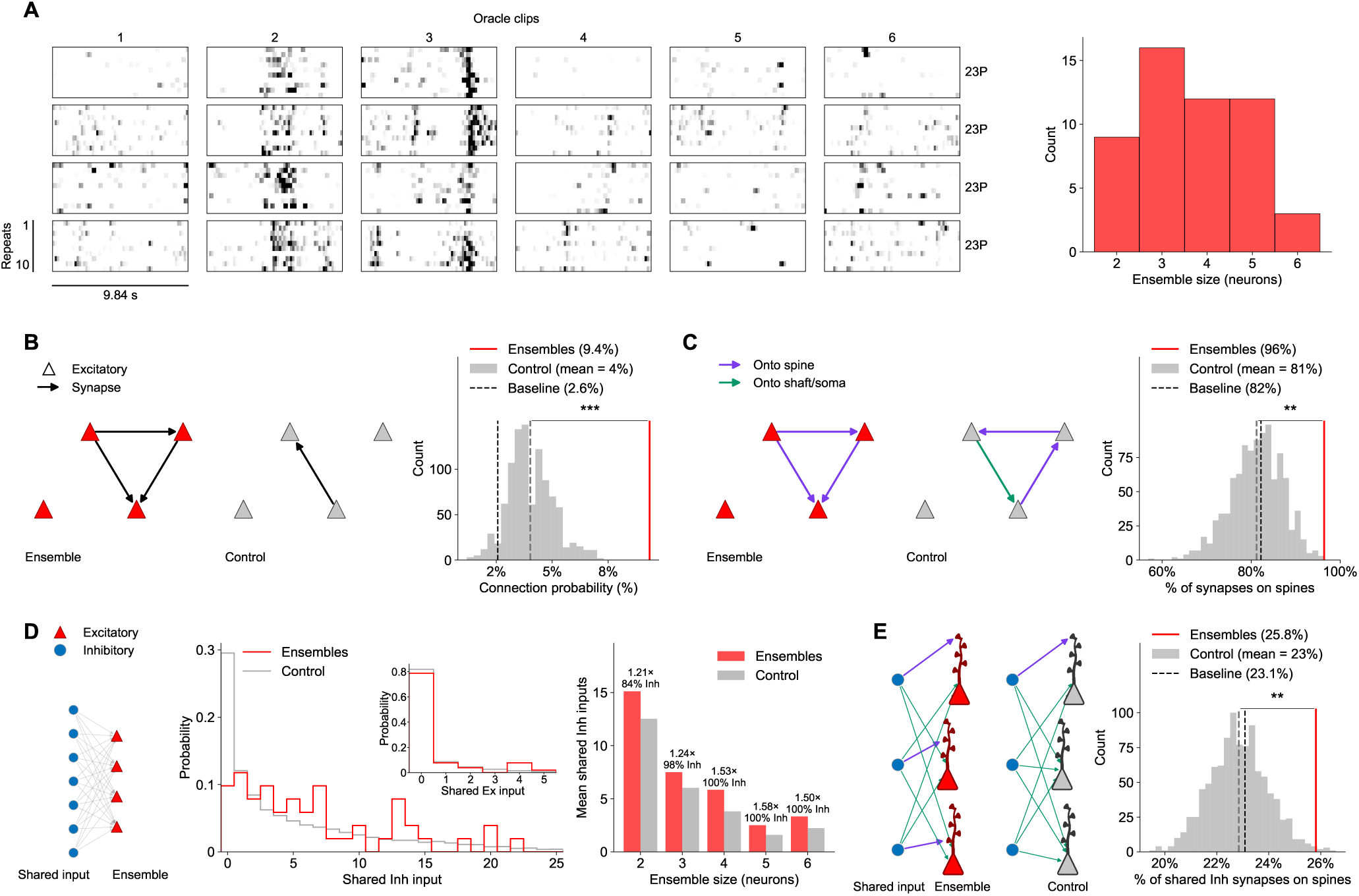
Neuronal ensembles are densely interconnected, spine-targeting sub-networks bound by shared local inhibition. **A.** *Left:* Raster plots of normalized deconvolved calcium activity for four excitatory (2/3P) ensemble members across six repeated oracle movie clips (10 repeats each). *Right:* Distribution of ensemble sizes across all identified ensembles (*n* = 52). **B.** Connection probability among ensemble members. *Left:* Schematic contrasting the total number of synapses among the neurons of a representative ensemble (red neurons with 3 synapses) with a size-matched control group (grey neurons, 1 synapse). *Right:* Connection probability across all ensembles (red line; 9.4%) against a bootstrap null distribution (grey; mean = 4% - grey dashed line; *p* = 0.001). Baseline of excitatory-to-excitatory (E→E) connection probability across the whole micro-column population is shown by the black dashed line; 2.3%. **C.** Spine targeting among ensemble members. *Left:* As in B, with synapses color-coded by ultrastructural target (purple -spine; green -shaft/soma). *Right:* Observed pooled percentage of internal synapses onto dendritic spines across all ensembles (red line; 96%) against a bootstrap null distribution (grey; mean = 81% - grey dashed line; *p* = 0.002), shown alongside the baseline E→E spine-targeting rate across the micro-column (black dashed line; 82%). **D.** Shared presynaptic input among ensemble members. *Left:* Schematic of the shared-input architecture; shared input is mostly composed of inhibitory neurons. *Middle:* Probability distribution of the number of shared inhibitory presynaptic partners across all ensembles (red) against a matched bootstrap control (grey); inset, same for shared excitatory partners. *Right:* Mean number of shared inhibitory partners for ensembles (red) and matched controls (grey), by ensemble size. Above each pair of bars: the observed/matched-null fold-change, and the percentage of the ensembles’ shared input at that size that is inhibitory. **E.** Spine preference of the shared inhibitory input from D. *Left:* Schematic illustrating spine (purple) versus shaft/soma (green) targeted by shared inhibitory synapses in ensemble (red neurons) and in control (grey neurons). *Right:* Observed pooled percentage of shared inhibitory synapses onto dendritic spines across all ensembles (red line; 25.8%) against a bootstrap null distribution using the same matched controls as in D (grey; mean = 23% - grey dashed line; *p* = 0.008), shown alongside the baseline inhibitory-to-excitatory (I→E) spine-targeting rate across the micro-column population (black dashed line; 23.1%).

We first asked whether members of a given ensemble are more likely to be directly wired to one another than expected by chance. Because nearby neurons are intrinsically more likely to be connected (Holmgren et al., 2003; Perin et al., 2011), we compared ensembles against random control groups matched not only for size but also for spatial compactness - how tightly clustered a group’s neurons are in physical space (see Methods); this compactness control is used throughout the rest of Figure 6. Ensemble members are connected to each other at a probability of 9.4%, 2.3-fold higher than matched controls (4%, *p* = 0.001), and well above the baseline excitatory-to-excitatory (E→E) connection rate across the micro-column population (2.3%; Fig. 6B). We next asked whether these internal connections preferentially target dendritic spines. Control groups here were matched not only for size and compactness (distance), as above, but also for the number of internal connections within the group - a density match, ensuring ensembles and controls are compared over a similar number of synapses (see Methods). Among the total 55 synapses formed between ensemble members, 96% targeted dendritic spines, compared to 81% in the matched controls (*p* = 0.002), and well above the E→E spine-targeting rate across the micro-column population (82%; Fig. 6C). The effect held over 209 internal synapses when ensembles were detected with a different algorithm (92% versus 75%, *p* = 0.001; Fig. S14C). Internal ensemble connectivity is therefore not only denser than expected, but also routed through spines.

We then extended the shared-input framework of Figure 4 to ask whether ensemble members are innervated by common upstream partners. Shared input is defined as the set of presynaptic neurons that innervate every member of a given ensemble - an all-to-all pattern of connectivity, analogous to a fully-connected layer in an artificial neural network (Fig. 6D, left; schematic as in Fig. 4A). Note that this is shared input from outside the ensemble and thus orthogonal to the increased connection probability within the ensemble described above. Control groups here were matched not only for size, compactness, and internal connectivity, as above, but also for each ensemble’s summed excitatory and inhibitory in-degree. An in-degree match was required because groups of high-in-degree neurons share more input than groups of low-in-degree neurons by chance alone. Summed across ensembles, ensemble members shared a total of 366 inhibitory presynaptic partners, against a matched-null expectation of 280 (Fig. 6D middle, *p* ≤ 0.001), an effect that strengthened with ensemble size, from 1.21-fold in ensembles of pairs of neurons to ∼1.5-fold in ensembles of four or more neurons (Fig. 6D, right). Shared presynaptic excitatory partners were comparatively rare: 29 observed against a matched-null expectation of 17 (Fig. 6D, middle inset, *p* = 0.017), and effectively absent in ensembles of four or more members.

At the same time, the mean shared input strength of ensemble members was 4.7% higher than the mean over a strong control (205.8 vs. 196.6, Fig. S15). While this difference did not reach statistical significance (*p* = 0.056), it did reach significance for ensembles from the alternative method (194.9 vs. 188.4; *p* = 0.002). This indicates that their shared excitatory input is shared *within* the ensemble, while the analysis in Fig. 6D only considers shared input from outside the ensemble. Note that our data source is limited to neuron pairs at relatively low distances and shared excitation from outside the ensemble could still exist at higher distances. In terms of balance, the composition of shared input shifted almost entirely toward inhibition as ensemble size increased, from 84% inhibitory in ensembles of pairs of neurons to 100% inhibitory in ensembles of four or more members (Fig. 6D, right).

Finally, we asked whether this convergent inhibitory input itself preferentially targets dendritic spines. Using the same matched controls as in Figure 6D, shared inhibitory synapses landed on spines at 25.8%, compared to 23% in matched controls (*p* = 0.008) and a baseline inhibitory-to-excitatory (I→E) spine-targeting rate of 23.1% across the micro-column population (Fig. 6E). This 25.8% is measured over 4,300 shared inhibitory synapses - by far the largest sample in the figure, compared to just 55 internal synapses underlying Figure 6C. Results for the alternative ensemble detection algorithm reproduced this effect over 9,986 shared inhibitory synapses (25.1% versus 22% in matched controls, *p* = 0.001; Fig. S14E).

Together, these results show that coactive ensembles are structurally distinct sub-networks already at the local micro-column level. They are more densely interconnected than chance, and these connections are made preferentially through spines. Their local presynaptic shared input is mostly composed of inhibitory neurons that themselves favor targeting spine. All of these results were reproduced using an independent ensemble-detection algorithm applied to the same recordings (Fig. S14), indicating that these wiring properties are features of coactive groups rather than of a particular detection method. We hypothesize that this dense, shared local inhibition provides the necessary structural constraint to shape, stabilize, and synchronize these small neuronal ensembles.

## Discussion

By combining large-scale MICrONS electron microscopy connectomics with matched *in vivo* calcium imaging, this study reveals a spine-based architecture that links cortical wiring to coordinated neuronal activity. Our findings revise the traditional view that dendritic spines serve to increase the number of synaptic contacts available to a neuron. Sparsely spiny inhibitory neurons receive nearly twice as many synapses, and at approximately twice the synaptic density, as spiny excitatory neurons, demonstrating that dense innervation does not require spines. Instead, excitatory and inhibitory neurons appear to follow distinct connectivity strategies: inhibitory neurons receive dense, predominantly shaft-based input, whereas excitatory neurons use spines to organize selective, diverse, and higher-order connectivity patterns (Fig. 1).

Consistent with the “connectivity and diversity” hypothesis (Yuste, 2011), spine density predicts both the number of incoming synapses and the diversity of presynaptic partners in excitatory and inhibitory neurons (Fig. 2). However, the role of spines extends beyond expanding afferent connectivity. On the output side, excitatory, but not inhibitory, axons increasingly target spines both with increasing axonal path distance from the soma (see also Loomba et al. (2022) and Schmidt et al. (2017)) and with increasing total synaptic output (Fig. 3). Highly divergent “broadcasting” excitatory neurons, therefore, expand their connectivity largely through spine-directed synapses. Moreover, neurons that receive inputs from such broad projectors, and consequently share more presynaptic partners with the surrounding network, also tend to possess extended axons that preferentially target spines. We term this relationship an input-to-spine/output-to-spine organization: a neuron’s incoming input is systematically related to the subcellular targets of its own outputs (Fig. 4).

This organization provides a subcellular basis for several established principles of cortical wiring. Common-neighbor motifs (Perin et al., 2011), long-tailed degree distributions (Piazza et al., 2025; Brunel, 2016), assortative connectivity (Ko et al., 2011; Cossell et al., 2015), and rich-club organization (Van Den Heuvel and Sporns, 2011) are generally described as graph-level properties. Our results suggest that these principles are partly implemented through the choice between spine and shaft targets. In particular, excitatory neurons receiving input from divergent, spine-targeting axons tend, themselves, to be divergent and spine-targeting, creating a spine-based, assortative, and highly interconnected subnetwork (Figs. 3 and 4). The persistence of these relationships across progressively smaller nested cortical volumes indicates that they are not specific to the dimensions of the original micro-column, although their statistical detectability naturally declines when very few neurons and synapses remain (Fig. 5).

These findings also suggest that some previously described cell-type targeting rules (Yuste, 2011; Douglas and Martin, 2004; Petreanu et al., 2009) may arise from a more fundamental preference for subcellular compartments. Long excitatory axons may preferentially innervate excitatory neurons partly because excitatory cells possess many more spines; correspondingly, inhibitory neurons with relatively high spine densities receive more excitatory partners than less-spiny inhibitory neurons (Fig. S1). Similarly, the shift along excitatory axons from targeting inhibitory neurons to targeting more distal excitatory neurons (Schmidt et al., 2017; Bodor et al., 2025) may partly reflect the progressive transition from shaft-directed to spine-directed synapses. Importantly, we note that the term “axonal spine preference” is descriptive rather than mechanistic, as it is typically spine outgrowth toward existing axonal boutons that establishes the spinous synapse (Yuste, 2023; Nägerl et al., 2007; Zito et al., 2009; Yuste and Bonhoeffer, 2004). One possibility is therefore that molecular matching between axonal boutons and nearby dendrites regulates spine formation, facilitating continued axonal growth among compatible neuronal populations. This remains speculative and will require direct experimental testing.

The *in vivo* functional data (Fig. 6) provide a direct test of whether the structural architecture described above is associated with coordinated activity. We identified neuronal ensembles solely from trial-to-trial co-fluctuations in Ca^2+^ activity during natural visual stimulation. We found that ensemble members are directly interconnected by excitatory synapses at more than twice the rate observed in spatially matched control groups, and that these synapses are routed almost exclusively onto dendritic spines (Fig. 6B–C). Thus, the anatomical signature of a functional ensemble is not simply that its neurons are connected, but, critically, where those connections terminate. These results indirectly support the prediction of (Ecker et al., 2024) that ensemble membership is boosted by dendritic clustering of excitatory connectivity via activation of dendritic nonlinearities and NMDA receptors, as these mechanisms are often associated with spines *in vivo* (Fu et al., 2012; Takahashi et al., 2012; Tazerart et al., 2020). This result also supports latent-ensemble or “iceberg” models (Yuste, 2026; Alejandre-García et al., 2022), in which coactive groups are embedded in pre-existing connectivity and can be recruited without requiring immediate large-scale synaptic rewiring.

A second and unexpected feature of these ensembles is their convergent local inhibition. Ensemble members share more presynaptic inhibitory partners than expected, after controlling for their spatial compactness, internal connectivity, and inhibitory in-degree. Shared excitatory presynaptic partners are comparatively rare and disappear almost entirely in the larger ensembles, whereas shared inhibition becomes increasingly dominant (Fig. 6D). This pattern suggests a division of labor across spatial scales. Because most excitatory synapses onto the local micro-column originate outside its reconstructed boundaries, long-range thalamic, top-down, or neighboring cortical inputs may provide the principal excitation that recruits an ensemble. The local circuit may then refine this drive through two complementary motifs: recurrent excitation routed onto dendritic spines binds ensemble members together, whereas convergent inhibition constrains, synchronizes, and stabilizes their collective activity. The enrichment of shared inhibitory contacts on spines (Fig. 6E) further suggests highly local gating of excitatory signals that promote ensemble co-activity, consistent with the powerful effects of spine-specific inhibition and dually innervated spines (Doron et al., 2017; Ofer et al., 2026).

Dendritic spines are particularly well suited to mediate this ensemble interaction. Their electrical and biochemical compartmentalization (Yuste and Denk, 1995; Cornejo et al., 2022; Yuste, 2023; Koch and Zador, 1993), high local input resistance, and recruitment of NMDA- and voltage-dependent nonlinearities (Segev and Rall, 1988; Rinzel and Rall, 1974), can amplify and selectively integrate excitatory inputs. At the same time, their capacity for synaptic plasticity (Yuste, 2011, 2023; Yuste and Bonhoeffer, 2001) allows individual connections to be modified without affecting neighboring inputs. Ensemble members therefore communicate not through generic excitatory synapses, but through specialized compartments capable of amplifying, isolating, gating, and independently modifying incoming signals. This architecture provides a plausible cellular substrate for attractor dynamics and pattern completion (Buzsáki, 2010; Hopfield, 1982): spine-targeted recurrent excitation could reinforce an emerging activity pattern, while shared, spatially precise inhibition could prevent uncontrolled recruitment and sharpen the temporal coordination of the selected ensemble. Figure 7 summarizes our proposed role of dendritic spines in organizing functional cortical ensemble.

**Figure 7.**
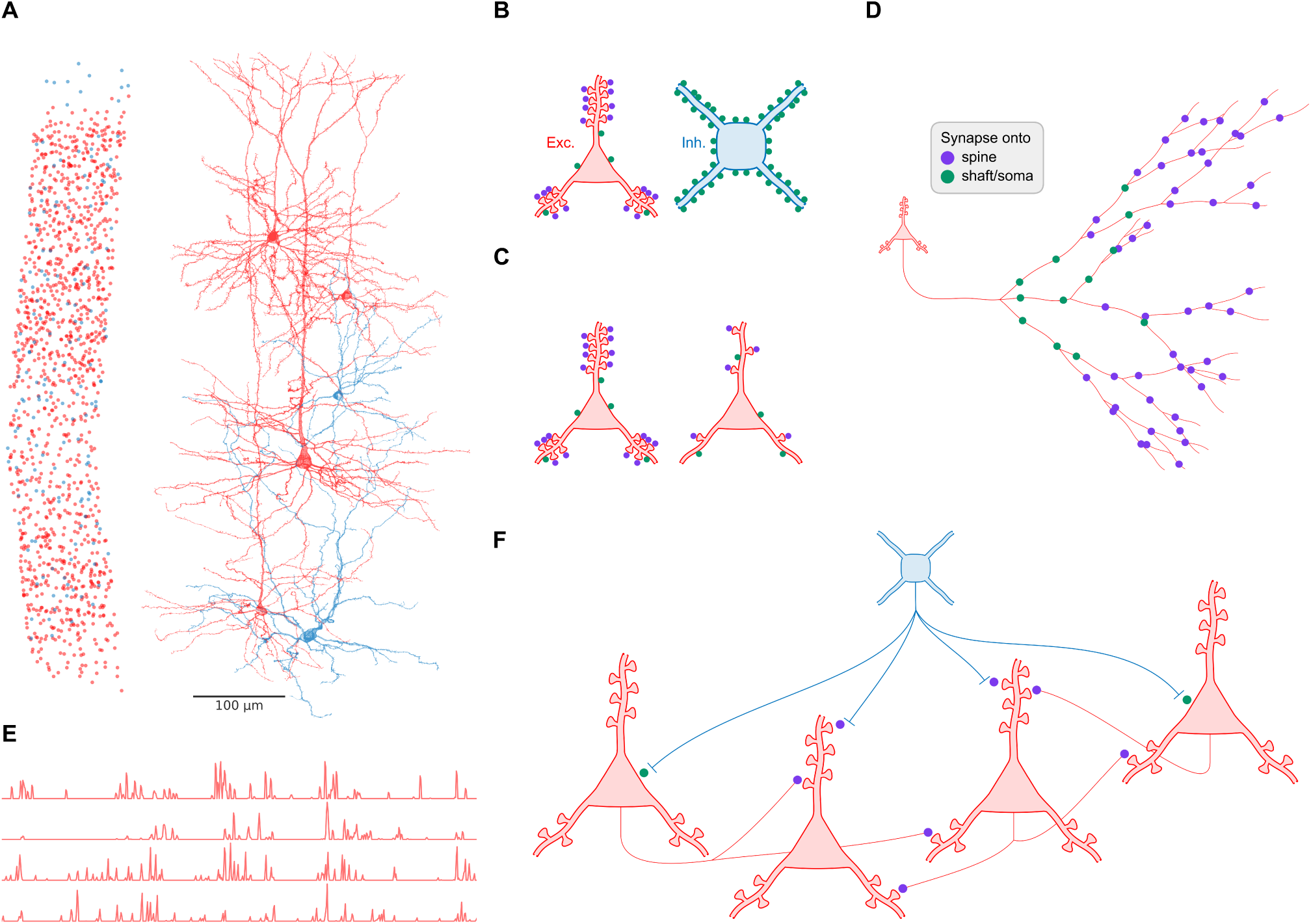
Proposed role of dendritic spines in organizing cortical ensembles. A. Connectivity. Reconstructed mouse V1 micro-column (as in Fig. 1A); excitatory neurons in red, inhibitory in blue. **B.** Inhibitory neurons receive denser and more synapses than excitatory neurons, mostly onto shafts; excitatory neurons are contacted mostly onto spines (Fig. 1). Purple, synapses onto spines; green, onto shaft/soma, throughout. **C.** More spines, more presynaptic partners (Fig. 2). **D.** Long-projecting excitatory axons prefer spines, a preference which increases with path distance from the soma (Fig. 3). **E. Activity.** Deconvolved calcium activity of excitatory neurons during natural movie viewing, from which ensembles were identified (Fig. 6A). **F. Functional ensemble.** Ensemble members are interconnected through spines and jointly innervated by inhibitory neurons that also prefer spines (Fig. 6) - a functional realization of the input-to-spine/output-to-spine motif of Figure 4.

These conclusions are limited by several constraints. The MICrONS micro-column contains only a fraction of the axons providing input to its neurons, and the long-range excitatory drive proposed above is therefore largely outside the reconstructed network. The structure–function relationships are observational and do not yet establish that spine targeting or shared inhibition causally generates ensemble co-activity. Moreover, the matched functional dataset samples only a subset of the reconstructed neurons (see Methods), and functional ensembles are necessarily small. Nevertheless, the principal results were reproduced using independent ensemble-detection methods (Fig. 6 and Fig. S14), supporting the interpretation that they reflect genuine circuit organization rather than being a property of a single analytical procedure.

In summary, as proposed before, dendritic spines indeed increase the connectivity space of presynaptic partners available to a neuron, but they do more. They organize which neurons exchange information and how functionally coactive neurons are bound into local subnetworks. Functionally defined ensembles are embedded in a distinctive architecture of dense recurrent spine-targeted excitation and convergent, predominantly inhibitory shared input that itself favors spines. Future work could test these principles in larger cortical volumes, determine the cellular rules governing spine targeting and formation, and use circuit simulations and causal manipulations to establish how the input-to-spine/output-to-spine motif shapes ensemble dynamics, learning, and pattern completion. Together, our findings identify dendritic spines as a structural interface through which cortical connectivity is transformed into coordinated neuronal activity.

## Methods

### MICrONS dataset annotation and structural tables

The MICrONS structural dataset consists of an electron microscopy volume of the visual cortex obtained from an adult mouse at postnatal day 87 (P87), corresponding primarily to the ‘minnie65’ reconstruction. All analyses utilized MICrONS dataset release version 1718 (March 7, 2026)^1^. To establish the spatial locations and baseline anatomical identities of neurons within our micro-column, we relied on the `nucleus_detection_v0` and `allen_v1_column_types_slanted_ref` tables (Schneider-Mizell et al., 2025). Synaptic connectivity between these neurons was retrieved from the `synapses_pni_2` table using the CAVEclient `synapse_query` method.

To analyze the specific ultrastructural landing sites of individual connections across the network, we utilized the `synapse_target_predictions_ssa_v2` table (Pedigo et al., 2026). This table provided compartmental predictions from which we derived the dendritic spine and dendritic shaft classifications used throughout the study. Finally, to track manual curation and ensure topological accuracy for specific validations - such as the extended network analysis presented in Figure S3 - we accessed the `proofreading_status_and_strategy` table^2^. For this analysis, which included neurons spanning outside the boundaries of the micro-column, we strictly limited our analysis scope to a highly curated subset of neurons that had undergone exhaustive manual correction (referred to by the Allen Institute as “axon_fully_extended”). Because this extended analysis included neurons outside our primary volume, their structural classifications were based on automated predictions from the Allen Institute’s `aibs_metamodel_celltypes_v661` table (Elabbady et al., 2025).

### Neuron selection and dataset filtering

Our initial target population comprised 1,351 neurons (1,188 excitatory and 163 inhibitory) within the micro-column. While the original reference dataset (Schneider-Mizell et al., 2025) defines 1,352 neurons for this volume, a single skeleton was excluded due to an unrecoverable download failure via CAVE. To ensure we were working with fully resolved structural and synaptic data, we applied a few standard inclusion criteria to this initial cohort. First, we filtered out neurons with zero dendritic or axonal length, as well as any cells missing compartmental mapping for their incoming spine-directed synapses. We also wanted to ensure reliable compartmental annotations for incoming connections, so we calculated a synaptic tagging ratio for each cell - defined as the sum of synapses successfully mapped to the soma, dendritic shafts, and dendritic spines, divided by the total number of incoming synapses. We retained the neurons where this mapped synapse ratio exceeded 0.9. Following these basic filtering steps, our final analytical dataset consisted of 1,298 neurons (1,139 excitatory and 159 inhibitory).

### Spine density trend analysis and binning

To robustly analyze the relationship between dendritic spine density and incoming connectivity metrics, mitigating the influence of extreme biological outliers, we grouped the neurons into specific density bins for the analyses presented in Figure 2 (and Fig. S2).

In our raw data, the measured spine density ranged from 0.1 to 1.8 spines/µm for excitatory neurons (Fig. 2C) and 0.04 to 1.2 spines/µm for inhibitory neurons (Fig. S1C). To create a consistent scale for our trendlines across these figures, we divided the data into custom bins with centers ranging from 0.25 to 1.25 spines/µm (using bin edges at 0.125, 0.375, 0.625, 0.875, 1.125, and 1.375). We then calculated both the average connectivity metric and the standard error of the mean (SEM) for the neurons within each bin. This approach provided reliable central tendencies to plot the data and allowed us to calculate the overall changes and multiplication factors across the analyzed trendline.

### Spine targeting distance along the axon

To quantify the spatial distribution of synaptic target preference, we analyzed the fraction of synapses targeting dendritic spines as a function of the axonal path distance from the presynaptic soma. To compute those distances from the presynaptic soma to individual synapses, pre-computed topological skeletal representations of the neurons were downloaded directly via the CAVEclient. Path lengths were then calculated for each synapse utilizing the built-in graph traversal logic within the MeshParty package (Dorkenwald et al., 2020). Outgoing synapses were sorted into discrete spatial bins (80–100 µm wide, depending on the specific figure). Within each bin, the spine-targeting fraction was computed as the number of synapses formed onto dendritic spines divided by the total number of outgoing synapses.

For population-level analyses (e.g., Fig. 3F, 3G, 3J, 3K, and Fig. S3), we utilized a neuron-weighted (“macro”) approach. The average spine fraction per bin was calculated as the mean of the individual axonal fractions within that bin, with population variance represented by the standard error of the mean (SEM). A synapse-weighted (“micro”) approach, which instead pools all synapses across the population prior to calculating the fraction, yielded highly similar results.

To precisely identify the transition from a shaft-biased to a spine-dominated targeting regime (Fig. S7), we determined a characteristic crossing distance - the path length where exactly 50% of synapses target spines. This threshold was mathematically derived via linear interpolation between the centers of the two consecutive bins bracketing the 0.5 fractional crossing: *x*_0_ +(*x*_1_ − *x*_0_) · [(0.5 − *y*_0_)*/*(*y*_1_ − *y*_0_)], where (*x*_0_*, y*_0_) and (*x*_1_*, y*_1_) represent the distance and spine fraction of the bins immediately before and after the threshold, respectively.

At the individual neuron level, crossing distances exhibited substantial diversity and a heavily right-skewed distribution. To capture this biological variance, we categorized single-cell behaviors into three distinct groups: (1) continuously spine-preferring neurons, which maintained a spine fraction above 0.5 from their first valid proximal bin (assigned a crossing distance of 0 µm); (2) transitioning neurons, which exhibited a clear crossing point within the reconstructed volume; and (3) shaft-preferring neurons that never crossed the 0.5 threshold along their traced length (assigned a crossing distance of ∞). Because the population-level curve (which averages the fractions per bin) naturally smooths over this heterogeneity, single-cell transition points (e.g., Fig. S7) are summarized using the median and interquartile range (IQR) of the finite crossing values, alongside the specific population percentages of the 0 µm and ∞ boundary cases.

### Shared input statistical analysis

To rigorously test the statistical significance of the shared input properties presented in Figure 4B, we employed a “blocked” configuration (CFG) null model. This degree-preserving null model was constructed by separately shuffling each of the four connectivity blocks (Excitatory-to-Excitatory (EE), Excitatory-to-Inhibitory (EI), Inhibitory-to-Excitatory (IE), and Inhibitory-to-Inhibitory (II)) within the adjacency matrix. By preserving the in- and out-degree distributions for each neuron within these specific sub-circuits, this method established a more constrained and robust baseline expectation compared to shuffling the entire network as a single piece.

To evaluate the multimodality of shared input strength distributions (Fig. 4B), we applied Hartigan’s Dip Test (Hartigan and Hartigan, 1985). This analysis was implemented using the diptest Python package^3^, which calculates the Dip statistic and associated *p*-value to test the null hypothesis of unimodality.

### Functional ensemble detection and analysis

To identify functionally coactive groups of neurons, we used the MICrONS functional dataset^4^, which pairs *in vivo* two-photon calcium imaging with the same structural volume analyzed above. Of the 16 available recording sessions (“scans”), 13 yielded at least one detected ensemble and were included in this analysis; the remaining three were excluded because no ensembles were detected (one scan) or because too few excitatory neurons were both recorded and structurally matched to support reliable detection (two scans). Structural connectivity for this analysis was drawn from all 1,188 excitatory neurons of the micro-column, rather than the 1,139-neuron subset used in Figures 1–5, since the additional structural quality filtering applied there was not relevant to this analysis. Of these, 476 excitatory neurons had matched functional recordings in at least one scan, with a median of 41 recorded excitatory neurons per scan (range 20–56).

Because each scan’s four imaging planes are sampled at different times within a frame (ms_delay, up to ∼159 ms across planes), we resampled every neuron’s trace onto the scan’s true frame-time clock - using its *ms_delay* and the scan’s recorded frame times - before any correlation-based detection step.

Ensemble detection was restricted to frames acquired during presentation of the repeated “oracle” movie clips: six distinct clips, 9.84 s each, shown for approximately ten repeats per scan, yielding 3,745 of approximately 40,000 total recorded frames (9.4%) available for detection. To isolate co-fluctuations not attributable to shared stimulus tuning, for each neuron and each clip we computed the trial-averaged deconvolved response across repeats and subtracted this average from every individual repeat, yielding a trial-residual trace. All ensemble detection was performed on these residual traces rather than on raw activity.

Candidate ensembles were identified by applying an independent component analysis (ICA)-based detection algorithm (Lopes-dos Santos et al., 2013) to the trial-residual activity of each scan. Briefly, the number of significant components, *K*, was determined from the eigenvalue spectrum of the neuron-by-neuron correlation matrix using a circular-shift-derived threshold, and FastICA was applied within the resulting *K*-dimensional subspace to extract independent sources. A neuron was assigned as a member of a component when its projection weight exceeded two standard deviations above the component’s weight distribution. Components in which member neurons exceeded 80% of the recorded population in that scan were discarded, as these reflect global, scan-wide fluctuations rather than a discrete subset of neurons. This procedure yielded 52 ensembles across the 13 included scans, ranging from 2 to 6 members, comprising 192 total ensemble memberships across 174 distinct neurons, a small number of which belonged to more than one ensemble. All main-text analyses (Fig. 6) were performed on this set of ensembles.

To verify that the results did not depend on how ensembles were defined, we repeated the entire analysis using a second, mechanistically unrelated detection algorithm (Carrillo-Reid et al., 2015; Herzog et al., 2021; Ecker et al., 2024). Briefly, the per-neuron-normalized trial-residual traces were summed across the population, and time bins whose summed activity exceeded a threshold derived from circularly shuffled surrogates were retained as candidate network events. These event bins were clustered by cosine similarity, with the number of clusters selected by the minimum of the Davies-Bouldin index over a scanned range, and each neuron was assigned to a cluster when its correlation with that cluster’s activity profile exceeded a shuffle-derived threshold, controlled at a 5% false discovery rate (Benjamini-Hochberg) across all neuron-by-cluster tests. This procedure yielded 181 ensembles across 14 scans, ranging from 2 to 15 members, comprising 743 total ensemble memberships across 401 distinct neurons. All downstream structural analyses and matched-control procedures were identical to those described in Figure 6, with the control pool correspondingly restricted to the 787 excitatory neurons not belonging to any detected ensemble. Results are reported in Figure S14. This algorithm assumes that activity arrives as discrete events, an assumption not well met by traces that are near-continuously active during movie presentation; it is therefore used here as an independent check rather than as a primary detector.

Structural connectivity of each ensemble was computed using the same micro-column connectome described above, so that every synapse could be resolved to its specific pre- and post-synaptic identity. Internal connection probability (Fig. 6B) was calculated by pooling, across all ensembles, the number of observed directed synaptic connections among all *N*(*N* − 1) ordered pairs of members and dividing by the pooled number of ordered pairs.

Internal spine targeting (Fig. 6C) was calculated over the same internal connections, as the pooled number of synapses landing on a dendritic spine divided by the pooled total number of internal synapses. Shared presynaptic input (Fig. 6D) was defined, for each ensemble, as the set of presynaptic neurons synapsing onto every member (i.e., the intersection of members’ presynaptic partner sets), computed separately for excitatory and inhibitory presynaptic identity and pooled (summed) across ensembles. Spine targeting of the shared inhibitory input (Fig. 6E) was calculated as the pooled fraction of synapses from these shared inhibitory presynaptic neurons onto ensemble members that landed on a spine, analogous to Panel C.

Statistical significance for each measure was assessed against matched random control groups rather than against the raw micro-column baseline (black dashed line across Fig. 6). Controls were drawn from a pool restricted to excitatory neurons in the micro-column that were not a member of any detected ensemble in any scan (i.e., excluding all 174 distinct ensemble neurons from the 1,188 excitatory neurons), so that the observed and null populations were fully disjoint. For each ensemble, up to 1,000 matched control groups of the same size were sampled from this pool. Beyond size, control groups were matched on up to three further properties, of increasing constraint:

- Compactness (Fig. 6, all panels). Group compactness - the mean pairwise soma-to-soma distance across all member pairs, was matched within the greater of 35% of the ensemble’s value or 15 µm.
- Internal connectivity (Fig. 6, panels C, D, E). Controls were additionally matched on the number of internal connected pre → post pairs, collapsing multiple contacts between the same pair, so that spine-targeting and shared-input comparisons were computed over comparably connected samples.
- In-degree (Fig. 6, panels D, E only). Controls were additionally matched on the ensemble’s summed in-degree, computed separately for excitatory and inhibitory presynaptic partners: for each member, we counted its number of distinct presynaptic partners of a given class, then summed this count across all members of the group. Each of the two resulting sums (excitatory, inhibitory) was matched within ±35%.

Null distributions were generated by a paired bootstrap. For each of 1,000 replicates, one matched control group was drawn per ensemble from its own pool of eligible matches; these control groups were then pooled exactly as the observed ensembles (i.e., numerators and denominators summed across all ensembles) and the resulting population-level statistic recomputed, yielding one null value per replicate and 1,000 replicates in total. Significance was assessed as a one-sided empirical *p*-value: *p* = (1 +number of null replicates ≥ observed)*/*(1 + 1,000), and fold-change is reported as the observed value divided by the mean of the null distribution. For a minority of ensembles - most often under the most constrained matching scheme used for Panels D and E, fewer than 1,000 distinct control groups satisfying all matching criteria could be found in the pool; in these cases, all control groups found were used, yielding a smaller null sample for that ensemble.

### Neuron identification in figures

The specific neurons visualized in the representative figures are identified using their `pt_root_id`s from the MICrONS dataset release version 1718. Reconstructions presented in Figures 1, 2 and 7 display only dendritic morphology, whereas the neurons in Figure 3 show both dendrites and axons. The selected neurons, along with their corresponding color coding used in the visual renderings, are detailed as follows:

- **Figure 1B**: Layer 2/3 pyramidal neuron (red), `pt_root_id` 864691135277186789; Layer 5 basket interneuron (blue), `pt_root_id` 864691135875777166.
- **Figure 2A**: Layer 4 pyramidal neuron (purple), `pt_root_id` 864691136618553563; Layer 6 IT pyramidal neuron (green), `pt_root_id` 864691135214643256.
- **Figure S1A:** Layer 2/3 Martinotti interneuron (purple), `pt_root_id` 864691135571356038; Layer 5 basket interneuron (green), `pt_root_id` 864691135875414414.
- **Figure 3A**: Layer 5 IT pyramidal neuron (purple), `pt_root_id` 864691135617152361.
- **Figure 3B**: Layer 6 CT pyramidal neuron (green), `pt_root_id` 864691136177632518.
- **Figure 3C**: Layer 4 basket interneuron (blue), `pt_root_id` 864691135562001633.
- **Figure 7A**: Layer 2/3 pyramidal neuron (red), `pt_root_id` 864691135277186789; Layer 4 pyramidal neu-

ron (red), `pt_root_id` 864691135503344477; Layer 5 PT pyramidal neuron (red), `pt_root_id` 864691135324963484; Layer 5 basket interneuron (blue), `pt_root_id` 864691135875777166; Layer 6 CT pyramidal neuron (red), `pt_root_id` 864691135503293250; Layer 6 basket interneuron (blue), `pt_root_id` 864691135106189901.

## Code availability

The code used in this study is available at *GitHub*: https://github.com/deangeckt/spines_connectivity_logic.

## Acknowledgements

We thank Ido Aizenbud, David Beniaguev, Toviah Moldwin, Jonathan Leibner, Daniela Yoeli, and Sapir Shapira, as well as all other members of Segev’s lab and members of the Allen Institute for Brain Science, for fruitful discussions and valuable feedback regarding this work. This work was supported by NINDS grant 1RM1NS132981-01, the Gatsby Charitable Foundation, the Drahi Family Foundation, and by a generous grant from the David and Inez Myers Foundation.

## Appendix. Normalized Common Incoming Neighbors and Incoming-Source Out-Degree Are Order-Equivalent

This document outlines the structural equivalence between the normalized sum of common incoming neighbors and the incoming-source out-degree in directed graphs.

### Setup

Let *G* = (*V, E*) be a directed finite graph with *n* = |*V* | nodes (where *n >* 1). Define the binary adjacency matrix *M* ∈ {0, 1}*^n^*^×*n*^ where *M_wu_* = 1 if and only if there is a directed edge from *w* to *u*.

For a node *v* ∈ *V*, let:

- *N* ^−^(*v*) = {*w* ∈ *V* : *w* → *v*} be the in-neighborhood of *v*.
- *d*^−^(*v*) = |*N* ^−^(*v*)| be the in-degree of *v*.
- *d*^+^(*w*) be the out-degree of a node *w*.

### Raw Quantities

Define two unnormalized quantities for each node *v*:

**Definition 1** (Total Common Incoming Neighbors). Summed over all other nodes:

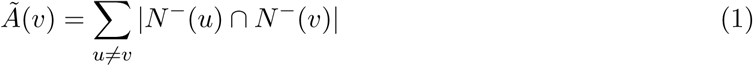

**Definition 2** (Total Out-Degree of Incoming Sources).

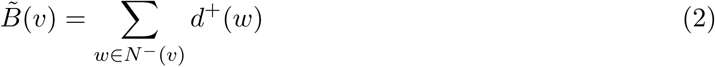

### Key Lemma

**Lemma 1.** For any node v ∈ *V,*

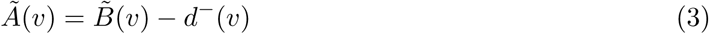

*Proof.* Express the intersection count using the adjacency matrix *M* :

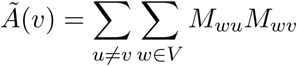

Swap the order of summation:

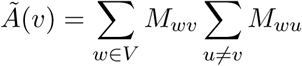

The inner sum counts the out-edges of *w*, excluding the edge to *v*. This holds true whether or not the graph contains self-loops:

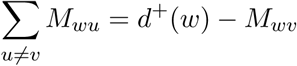

Substituting this back into the equation:

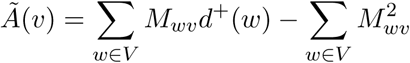

Because *M* is a binary matrix, *M* ^2^ = *M_wv_*. Therefore, the second sum is exactly the in-degree *d*^−^(*v*), yielding:

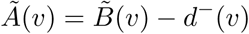

### Normalized Metrics and Their Relationship

For any node *v* where *d*^−^(*v*) *>* 0, we define the normalized metrics:

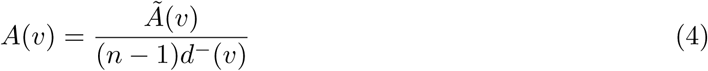

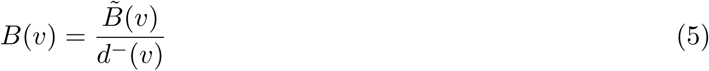

Dividing both sides of Lemma 1 by (*n* − 1)*d*^−^(*v*) yields:

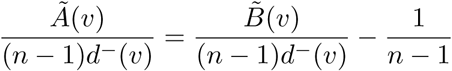

Which simplifies to the exact identity:

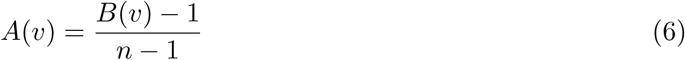

### Order Equivalence

**Corollary 1**. Because the transformation A(*v*) = *<u>^B^</u>*<u>^(*v*)−1^</u> *is strictly increasing with respect to B*(*v*)*, the following holds for any nodes u, v* ∈ *V (provided both have an in-degree greater than zero):*

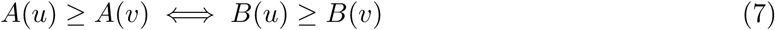

## Supplementary Figures

**Figure S1.**
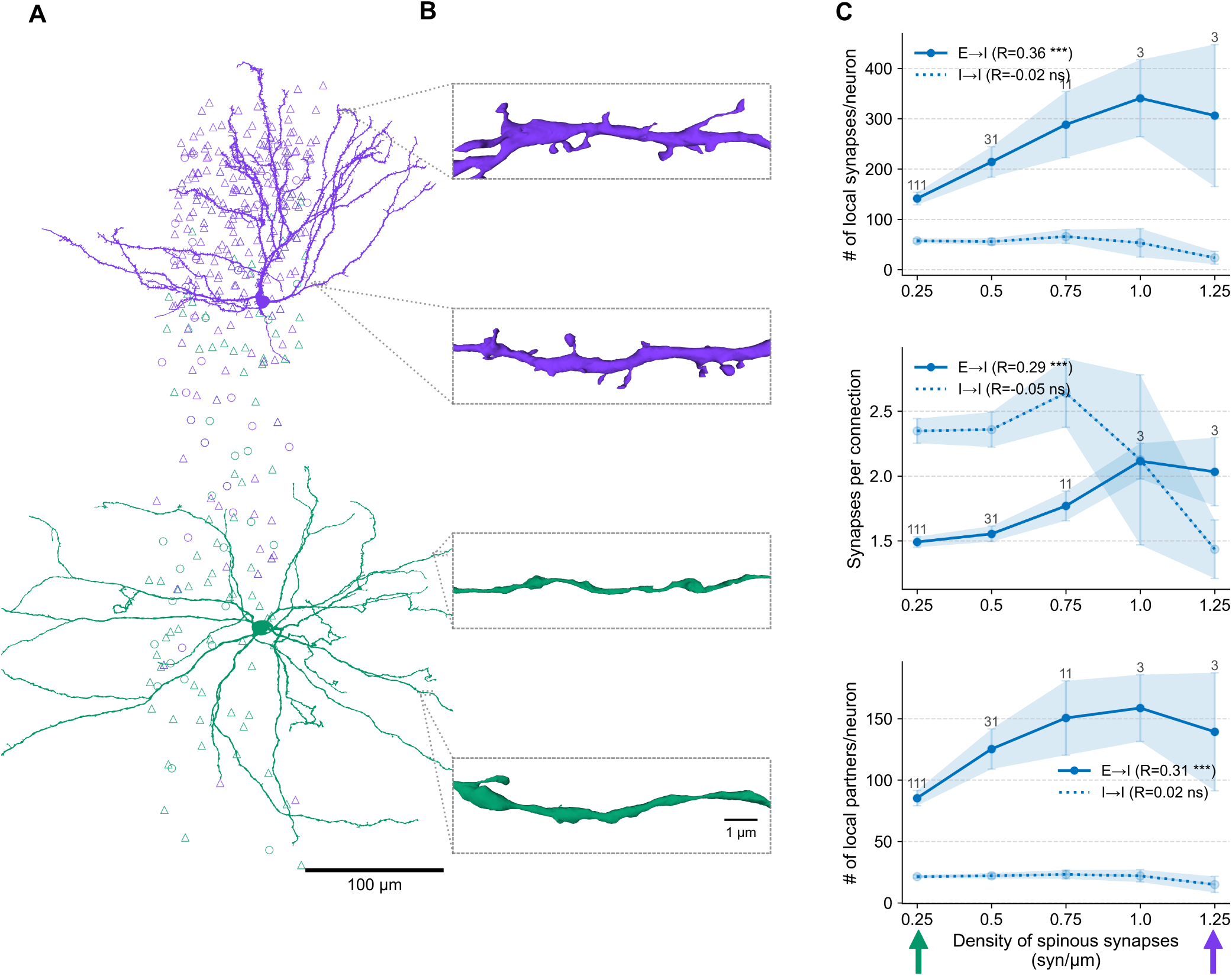
Spine density in inhibitory neurons specifically correlates with excitatory input richness and diversity. **A.** Example reconstruction of a layer 2/3 Martinotti interneuron (purple) and a layer 5 basket interneuron (green); only dendrites are shown. **B.** Representative dendritic segments from neurons shown in A. **C.** Equivalent analysis as in Figure 2, but for the inhibitory neuronal population.

**Figure S2.**
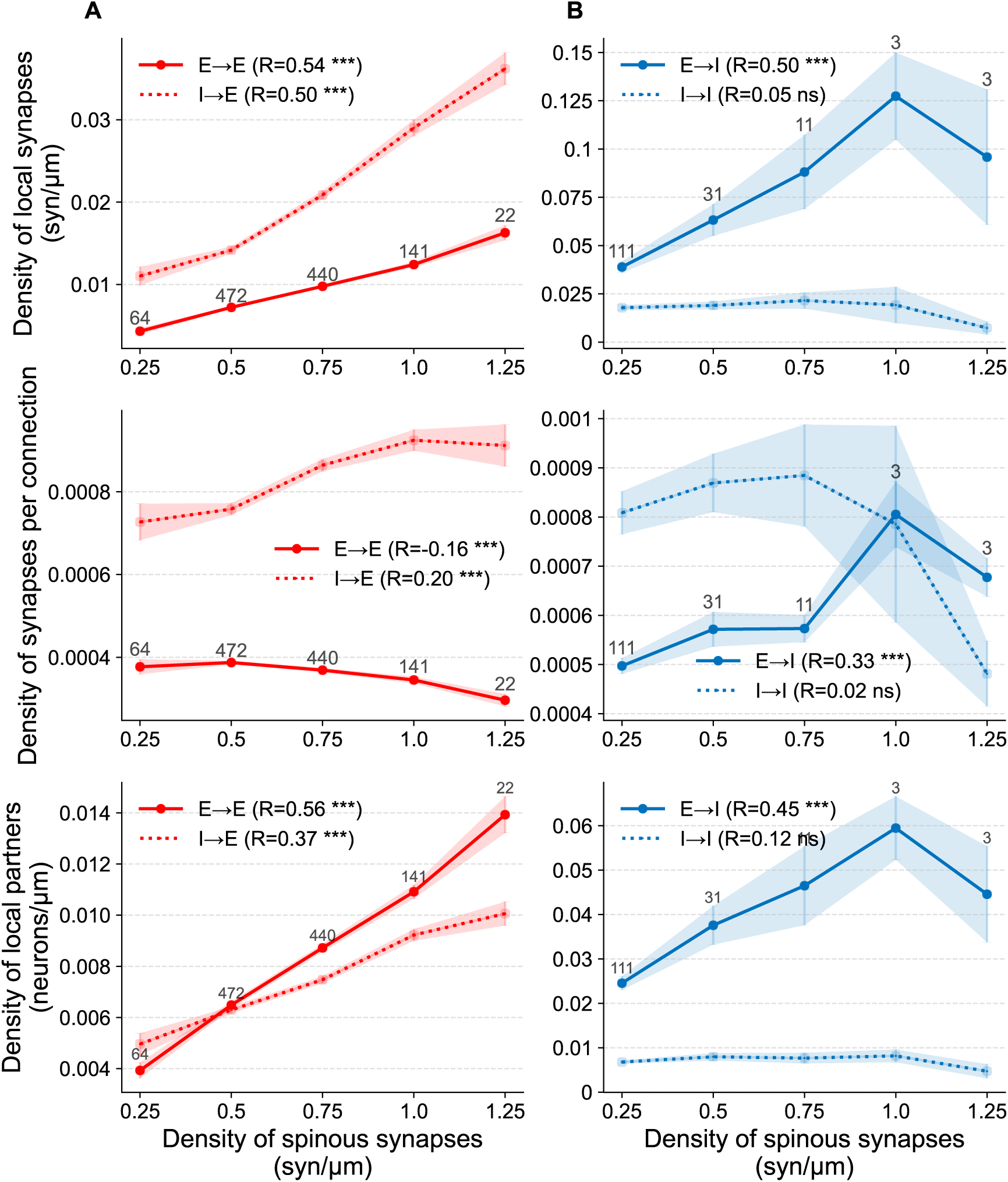
Synaptic input richness and diversity normalized by dendrite length. **A.** Normalized metrics versus total spine density for excitatory neurons. Unlike Figure 2C, all *y*-axes values are normalized by the total dendrite length of the (post-synaptic) excitatory neuron. *Top:* Density of local synaptic inputs as a function of spine density. *Middle:* Density of synapses per connection as a function of spine density. *Bottom:* Density of distinct presynaptic partners as a function of spine density. Solid lines: excitatory-to-excitatory synapses; dashed: inhibitory-to-excitatory synapses. **B.** As in A, but for inhibitory neurons (corresponding to Fig. S1C).

**Figure S3.**
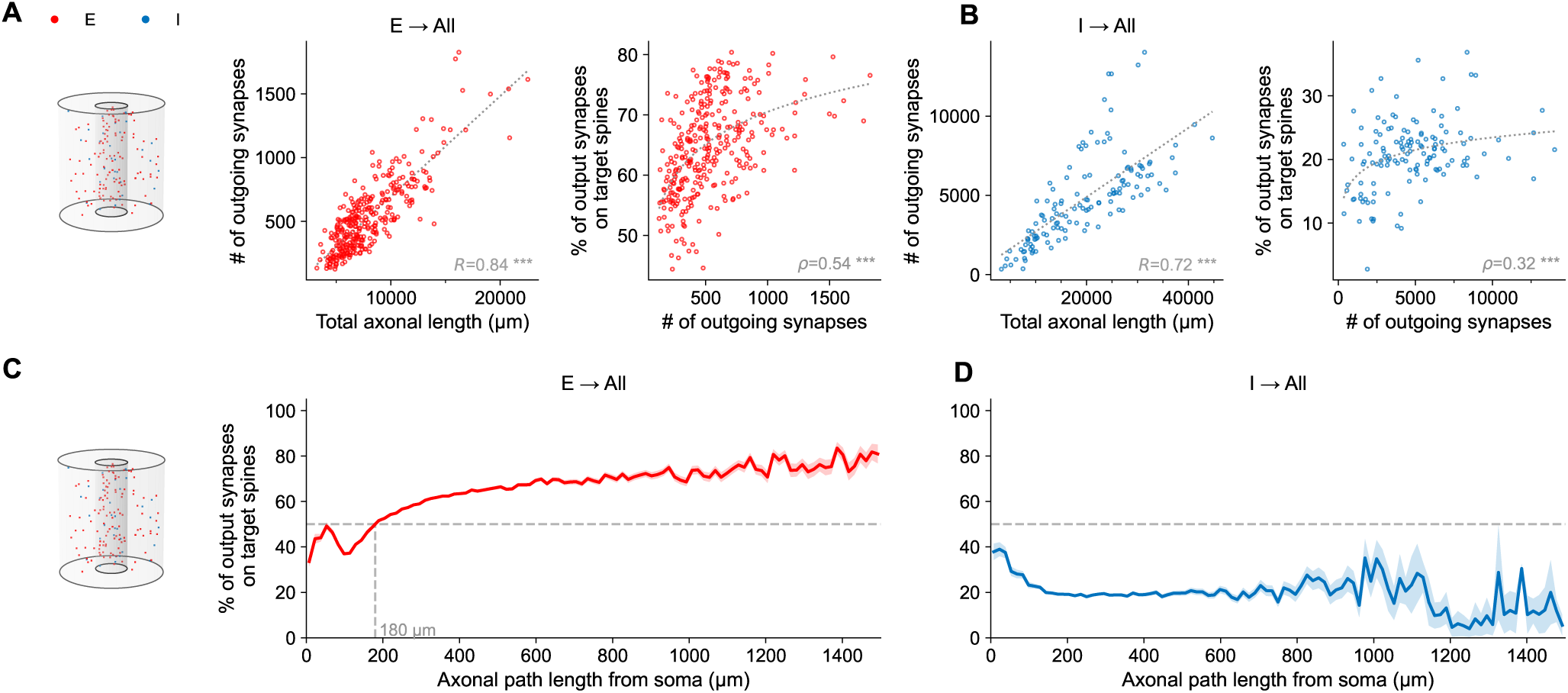
Distance-dependent spine targeting is robust across the most carefully proof-read reconstructed axons. Analyses as in Figure 3 but on a strict subset of neurons with the highest level of manual proofreading of their axons, consisting of 272 excitatory and 7 inhibitory neurons from within the micro-column, plus 42 excitatory and 126 inhibitory neurons from outside the micro-column. **A.** Analysis of synapses made by excitatory axons originating both inside and outside the micro-column onto the whole reconstructed circuit (schematic at left). Neurons’ cell type outside the micro-column are classified based on the Allen Institute *aibs_metamodel_celltypes_v661* table (Elabbady et al., 2025, and see Methods). *Left:* Number of excitatory axonal synapses correlates linearly with total axonal length. *Right:* Percentage of excitatory synapses onto dendritic spines increase as a function of the total number of outgoing synapses also in this highly curated group of axons. **B.** As in A but for the inhibitory axons. *Left:* Number of inhibitory synapses is linearly correlated with axonal length. *Right:* Percentage of synapses onto spines shows weaker dependence on the number of outgoing synapses. **C.** Spine targeting as a function of path distance along excitatory axons. The monotonic increase and spatial transition from a mixed regime near the soma to a spine-dominated regime at larger distances remain clearly evident. **D.** Same as in C for inhibitory axons. Spine targeting remains low and largely independent of path distance along the inhibitory axon.

**Figure S4.**
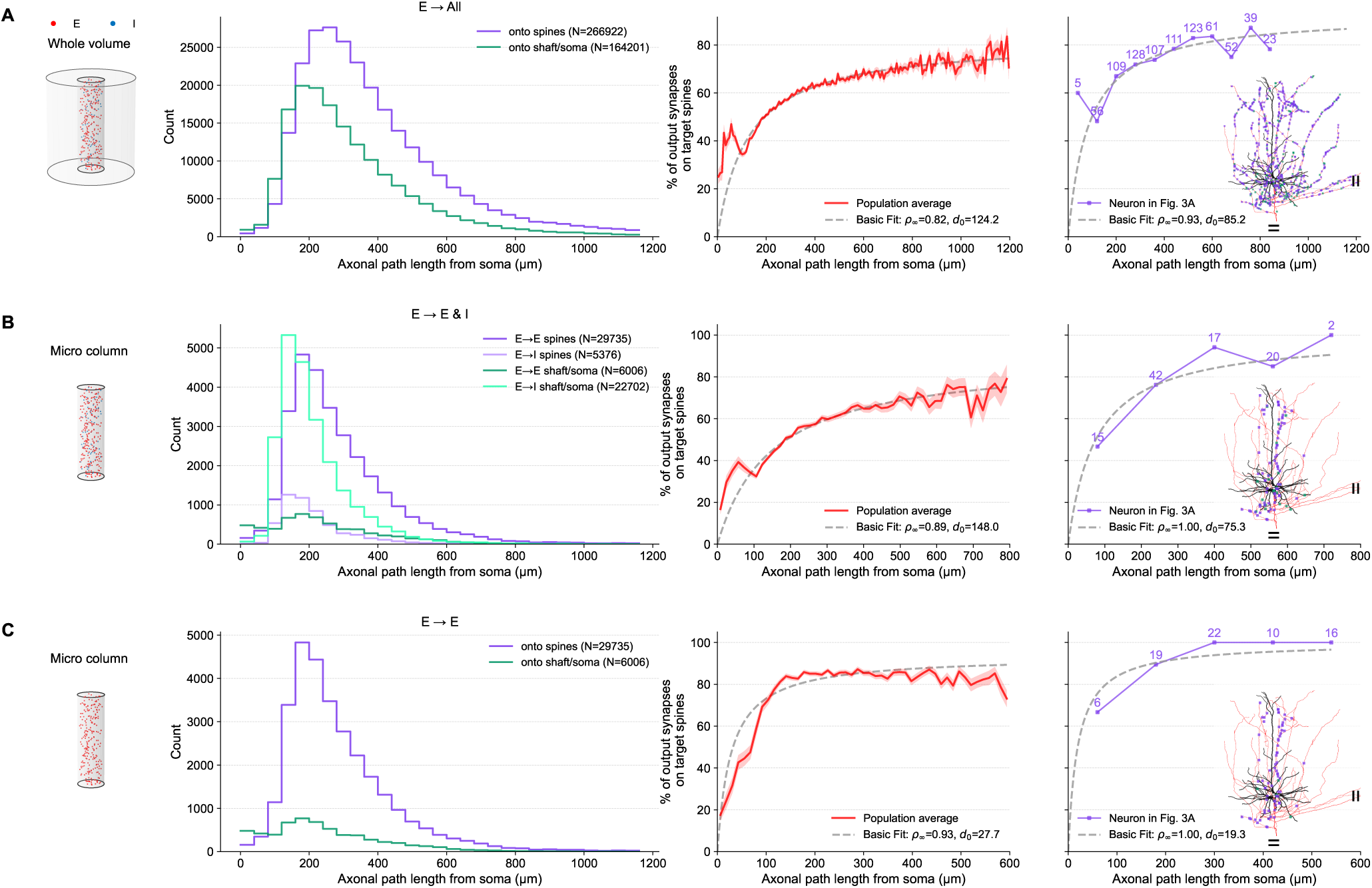
A minimal model of spine targeting preference as a function of axonal path length. **A.** Whole-volume analysis of all outgoing excitatory synapses. *Left:* Distribution of the number of synapses along the axonal path length, categorized by their target (purple for spines, green for shafts). *Middle:* Population-level target preference showing the percentage of synapses targeting spines (in red). The fitted model using Eq. (1) is shown by the dashed line; *ρ*_∞_ = 0.82, *d*_0_ = 124. *Right:* As in Middle but for a single representative L5-IT pyramidal neuron, also shown in Fig. 3A (number of synapses per bin are indicated above); fitted model is shown by the dashed line (*ρ*_∞_ = 0.93, *d*_0_ = 85.2). **B.** Analysis of local excitatory connections restricted to the micro-column. *Left:* Synapse distributions for four local targeting categories: onto spines of excitatory (E→E) and inhibitory (E→I) neurons (dark and light purple, respectively), and onto shafts or somas of E and I neurons (dark and light green, respectively). *Middle:* Population-level model fit for all targets within the local micro-column (*ρ*_∞_ = 0.89, *d*_0_ = 148). *Right:* Single-neuron fit restricted to these local micro-column targets (*ρ*_∞_ = 1, *d*_0_ = 75.3). **C.** Local E→E subnetwork analysis within the micro-column. *Left:* Distribution of synapses restricted exclusively to excitatory-to-excitatory connections, targeting spines (purple) versus shafts/somas (green). *Middle:* Population-level model fit for the E→E subnetwork (*ρ*_∞_ = 0.93, *d*_0_ = 27.7). *Right:* Single-neuron model fit for local E→E targets (*ρ*_∞_ = 1, *d*_0_ = 19.3).

**Figure S5.**
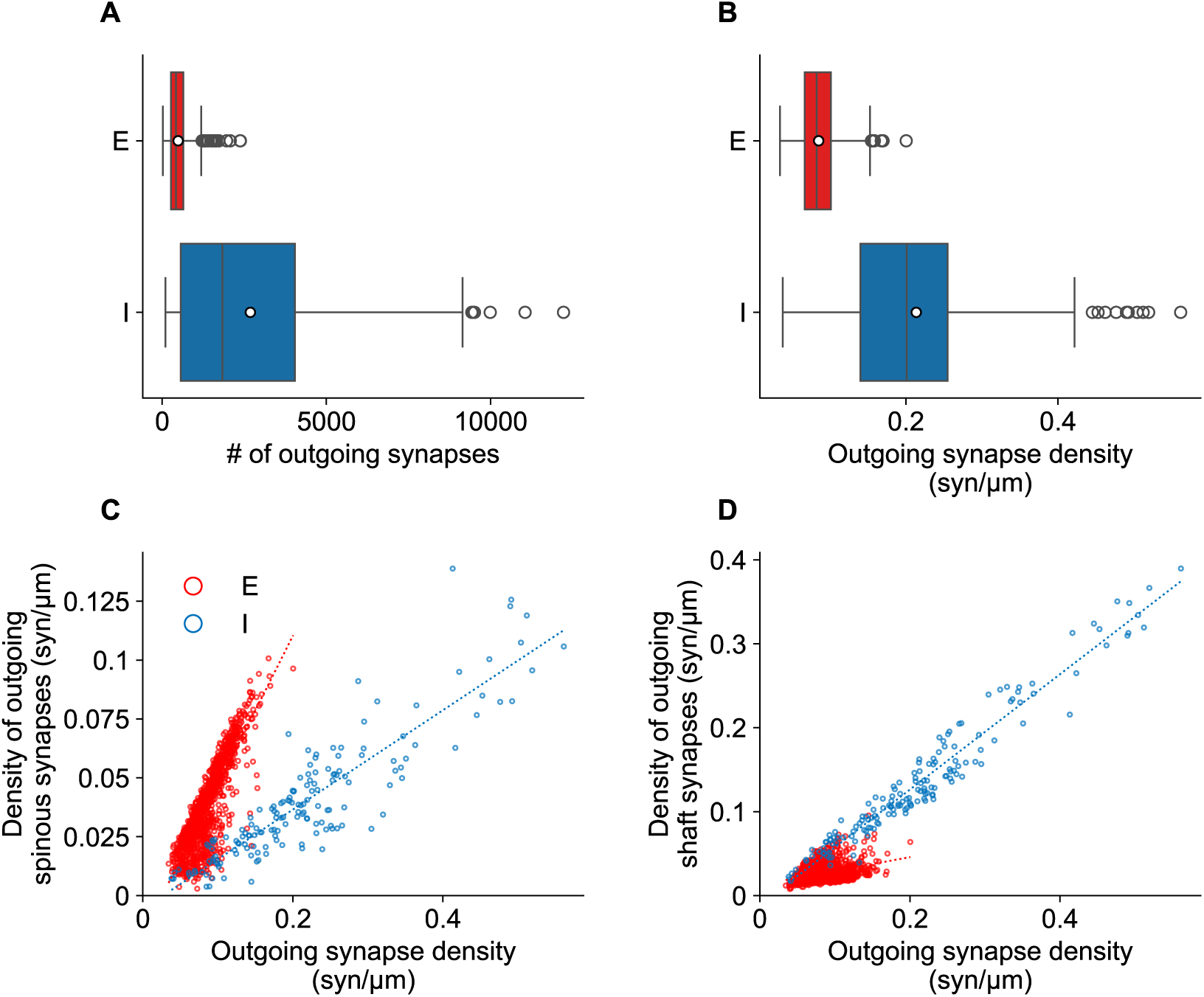
Divergent axonal targeting strategies: Excitatory subtypes exhibit diverse target preferences, while inhibitory neurons uniformly target dendritic shafts. **A–B.** Distribution of outgoing synaptic outputs for excitatory (red) and inhibitory (blue) neurons. **A.** The total number of outgoing synapses per neuron; **B.** Density of outgoing synapses. In both panels, the box edges represent the 25th and 75th percentiles, the vertical line denotes the median, and the white circle indicates the mean. Whiskers extend to 1.5 times the interquartile range. Mean ± SD values: **A**, excitatory: 488 ± 319 vs. inhibitory: 2,686 ± 2,623; **B**, excitatory: 0.09 ± 0.02 vs. inhibitory: 0.21 ± 0.11. **C.** Density of outgoing spinous synapses as a function of total outgoing synapse density for excitatory and inhibitory neurons. Linear regressions are shown by the dotted lines (excitatory: slope = 0.21, *R*^2^ = 0.80, *p <* 0.001; inhibitory: slope = 0.17, *R*^2^ = 0.16, *p <* 0.001). As a population, excitatory neurons show a robust correlation between total number of output synapses and number of spine targeted. **D.** Density of outgoing shaft synapses as a function of total outgoing density. Linear regressions are shown by dotted lines (excitatory: slope = 0.63, *R*^2^ = 0.71, *p <* 0.001; inhibitory: slope = 0.69, *R*^2^ = 0.95, *p <* 0.001). The scaling of inhibitory output density is driven almost entirely by shaft-targeting synapses.

**Figure S6.**
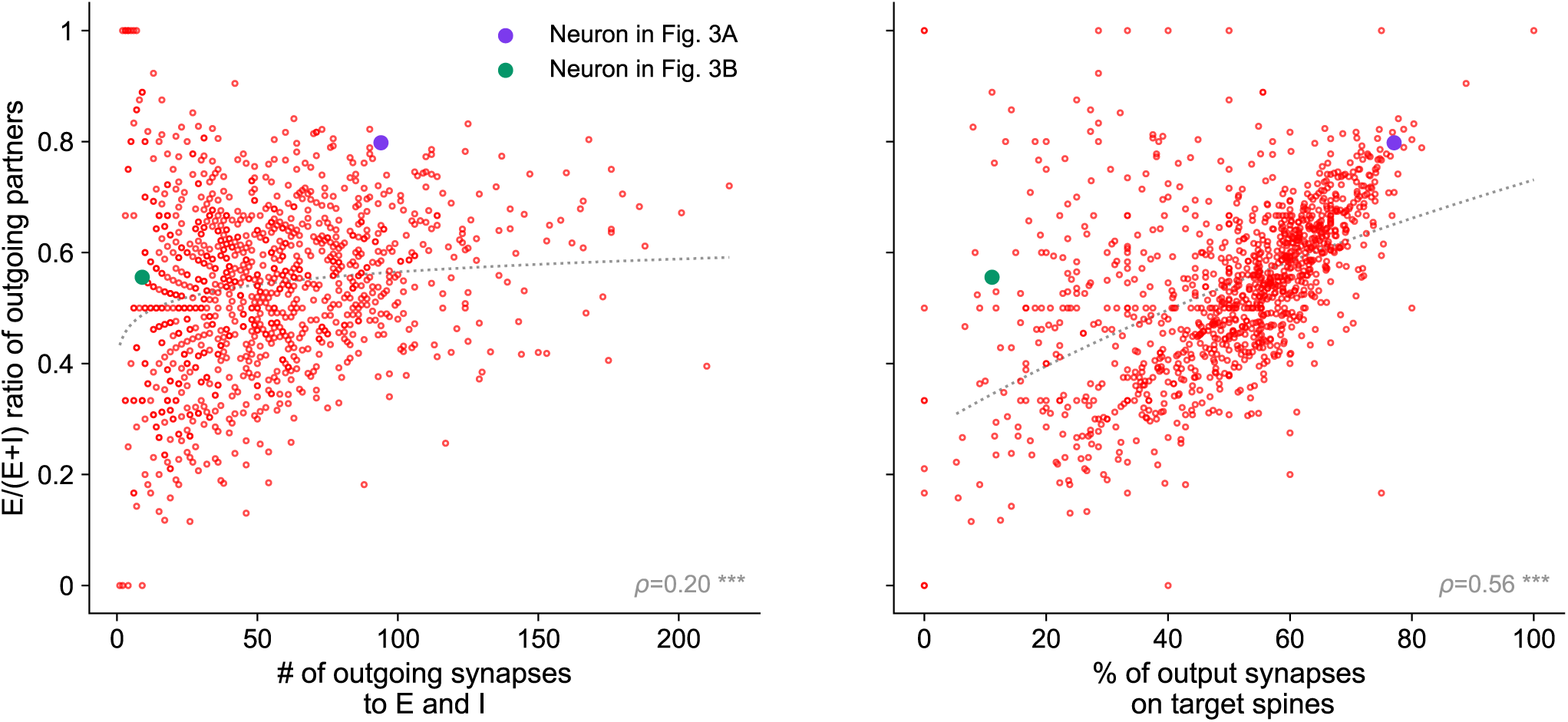
Spine preference is a stronger predictor of excitatory partner selection than total synaptic output. Analysis of local connectivity restricted to the micro-column, evaluating both excitatory and inhibitory targets for presynaptic excitatory neurons (red circles). The *y*-axis in both panels represents the target cell-type preference, calculated as the ratio of unique excitatory partner neurons to total local partner neurons [E / (E+I)]. *Left:* The ratio of excitatory partners shows only a weak correlation with the total number of outgoing synapses to both E and I targets (Spearman *ρ* = 0.20, *p <* 0.001). *Right:* The neuron’s overall preference for targeting dendritic spines (the percentage of output synapses onto local spines) is a much stronger predictor for its preference to connect with excitatory partners (Spearman *ρ* = 0.56, *p <* 0.001). Highlighted neurons correspond to the representative L5-IT (purple, neuron in Figure 3A) and L6-CT (green, neuron in Figure 3B) cells shown in Figure 3. This indicates that the tendency to target excitatory neurons is tightly coupled to an underlying subcellular preference for dendritic spines.

**Figure S7.**
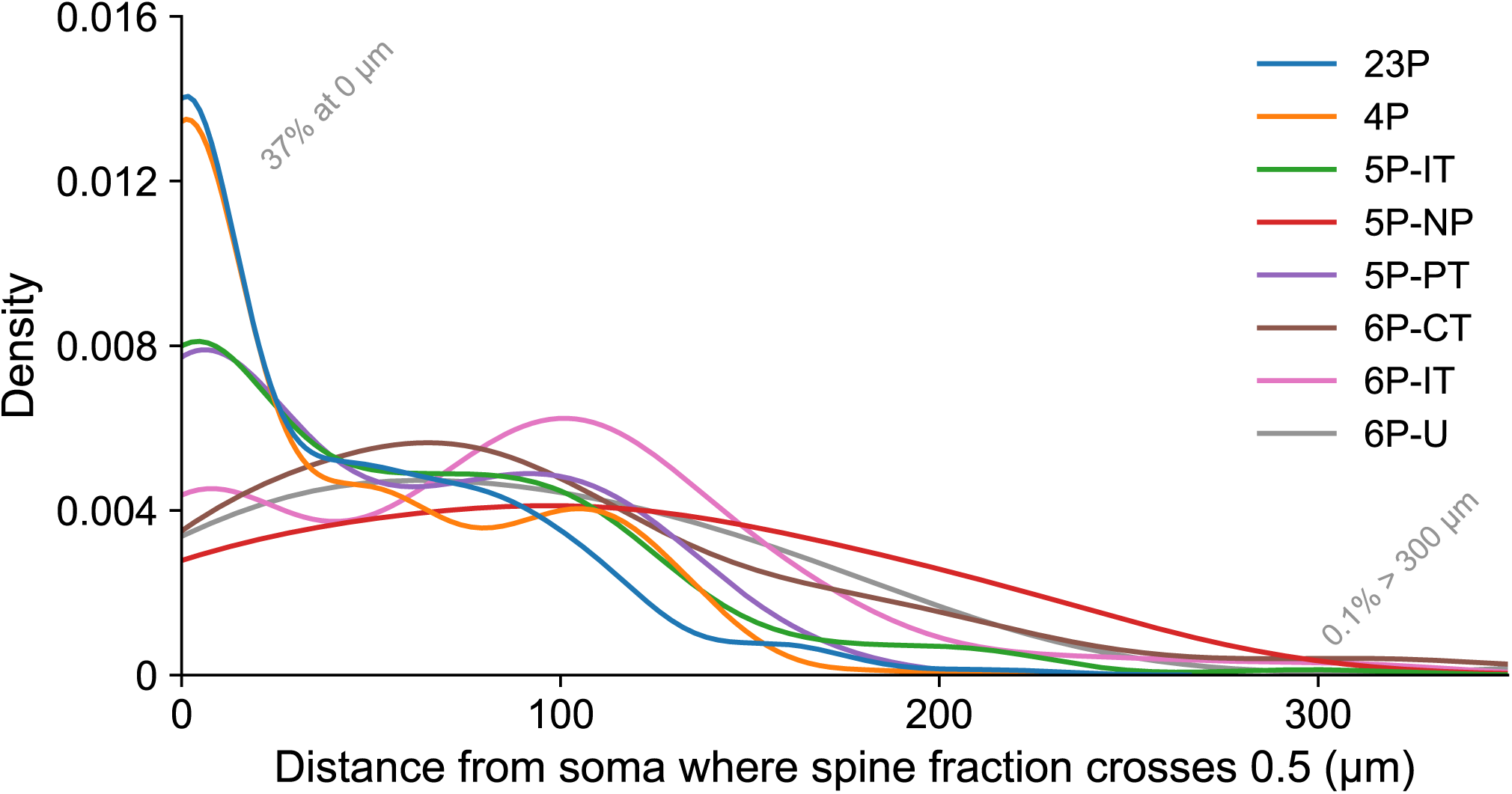
Distribution of axonal path distance from soma where spine targeting fraction crosses 50%. Kernel density estimate (KDE) of the distance from the soma where the spine-targeting fraction crosses the 50% threshold for individual excitatory neurons. This analysis incorporates all outgoing synapses from the E population within the 1 mm^3^ volume, serving as an extension to the population-level analysis in Figure 3F. Distributions are color-coded by cell type, with each normalized independently. Note that a substantial subset of neurons (37%) are “always above” the threshold (i.e., their crossing distance is 0 µm), preferring spines throughout, while a negligible fraction (0.1%) never reaches the 50% threshold within the volume, i.e., remained shaft-preferring across their entire reconstructed length.

**Figure S8.**
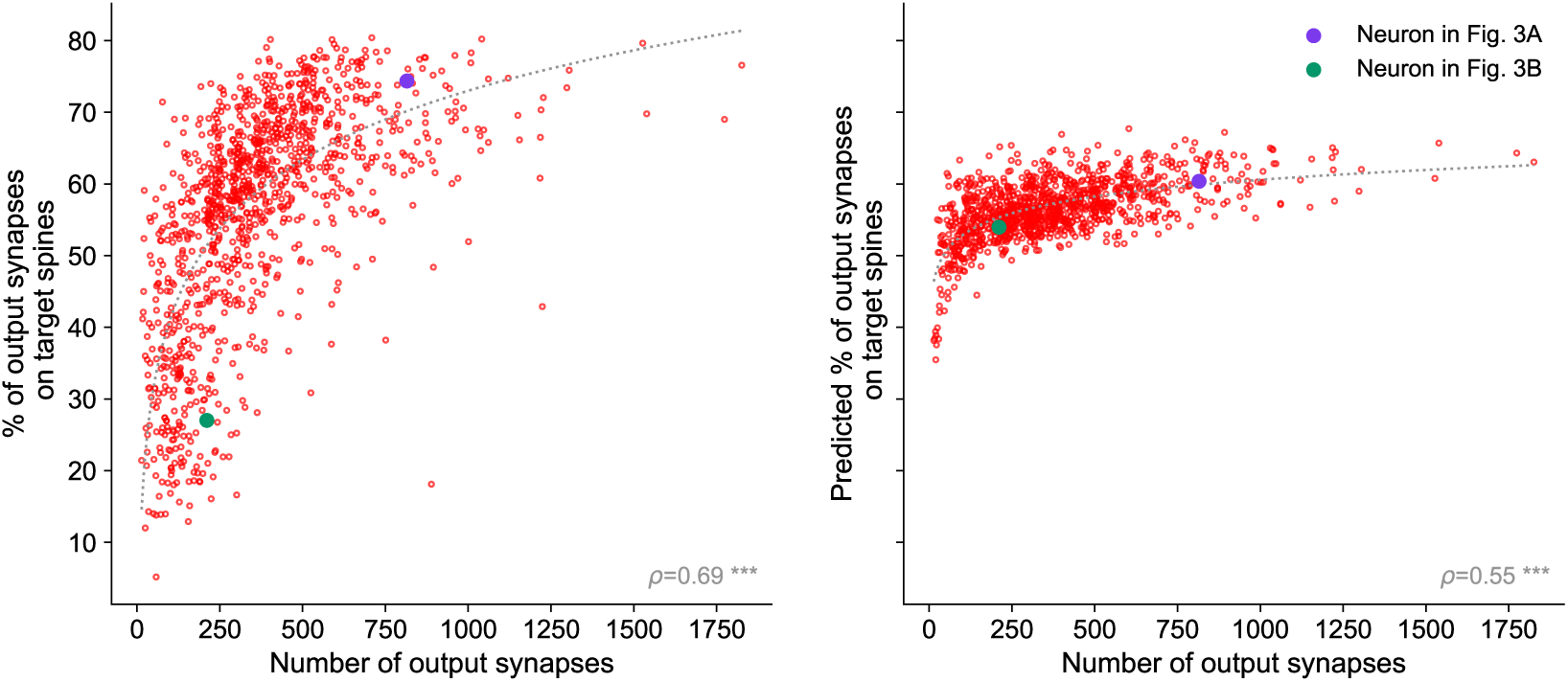
Spine preference variability cannot be fully explained by distance-based axonal mass distribution. Comparison of observed versus predicted spine preference for individual excitatory neurons. *Left:* Observed global spine preference as a function of total outgoing synapses per neuron (as in Figure 3D). *Right:* Predicted spine preference for the same neurons. Predictions rely solely on spatial distribution, generated by passing the empirical path length of every outgoing synapse through the population-fitted minimal model (*ρ*_∞_ = 0.82, *d*_0_ = 124.2 µm). While this distance-based model preserves a baseline upward trend (Spearman *ρ* = 0.55, *p <* 0.001), it fails to reproduce the wide empirical variance and the full magnitude of spine preference seen on the left.

**Figure S9.**
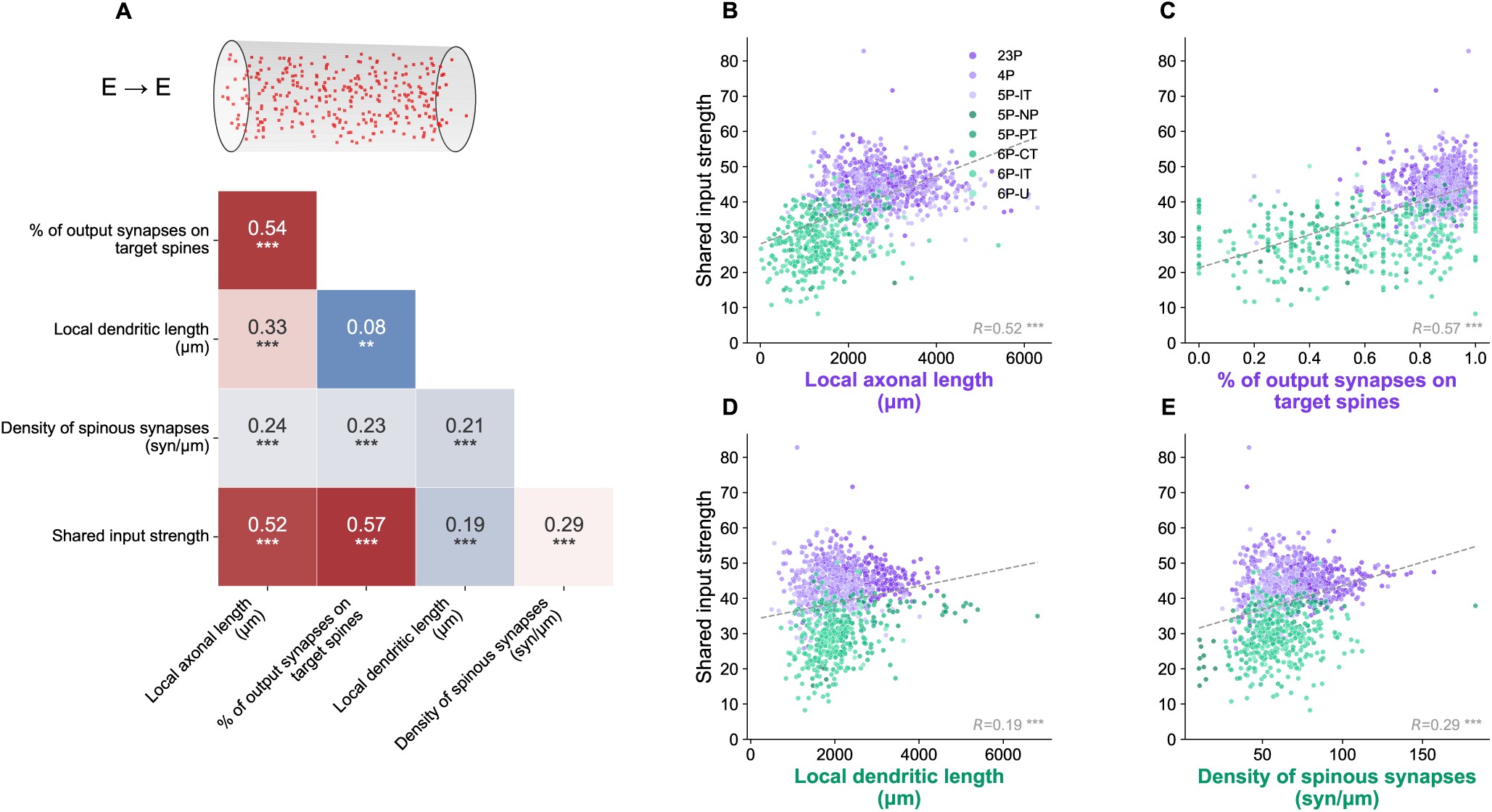
Feature correlations determining shared input strength within the excitatory sub-network (E-E connectivity). **A.** *Top:* Schematic representation indicating that this analysis is restricted to the excitatory-to-excitatory (E-E) connections within the micro-column. *Bottom:* A 5 × 5 Pearson correlation matrix displaying the relationships between shared input strength and four distinct structural features. For clarity, only the lower triangular portion of the matrix is shown, omitting the mirrored upper half and the diagonal of ones. **B–E.** Scatter plots detailing the relationship between a neuron’s shared input strength (*y*-axis) and individual network features (*x*-axis) for the E-E sub-network. Neurons are color-coded based on their cell type, with high-common-input neurons (2/3P, 4P, 5P-IT) in purple and the remaining neurons in green (classification detailed in main text). Panels B and C display correlations with “output” features (Pearson *R* = 0.52, *p <* 0.001 and *R* = 0.57, *p <* 0.001, respectively), which notably exhibit stronger correlations, while panels D and E display correlations with “input” features (Pearson *R* = 0.19, *p <* 0.001 and *R* = 0.29, *p <* 0.001, respectively) with the shared input strength.

**Figure S10.**
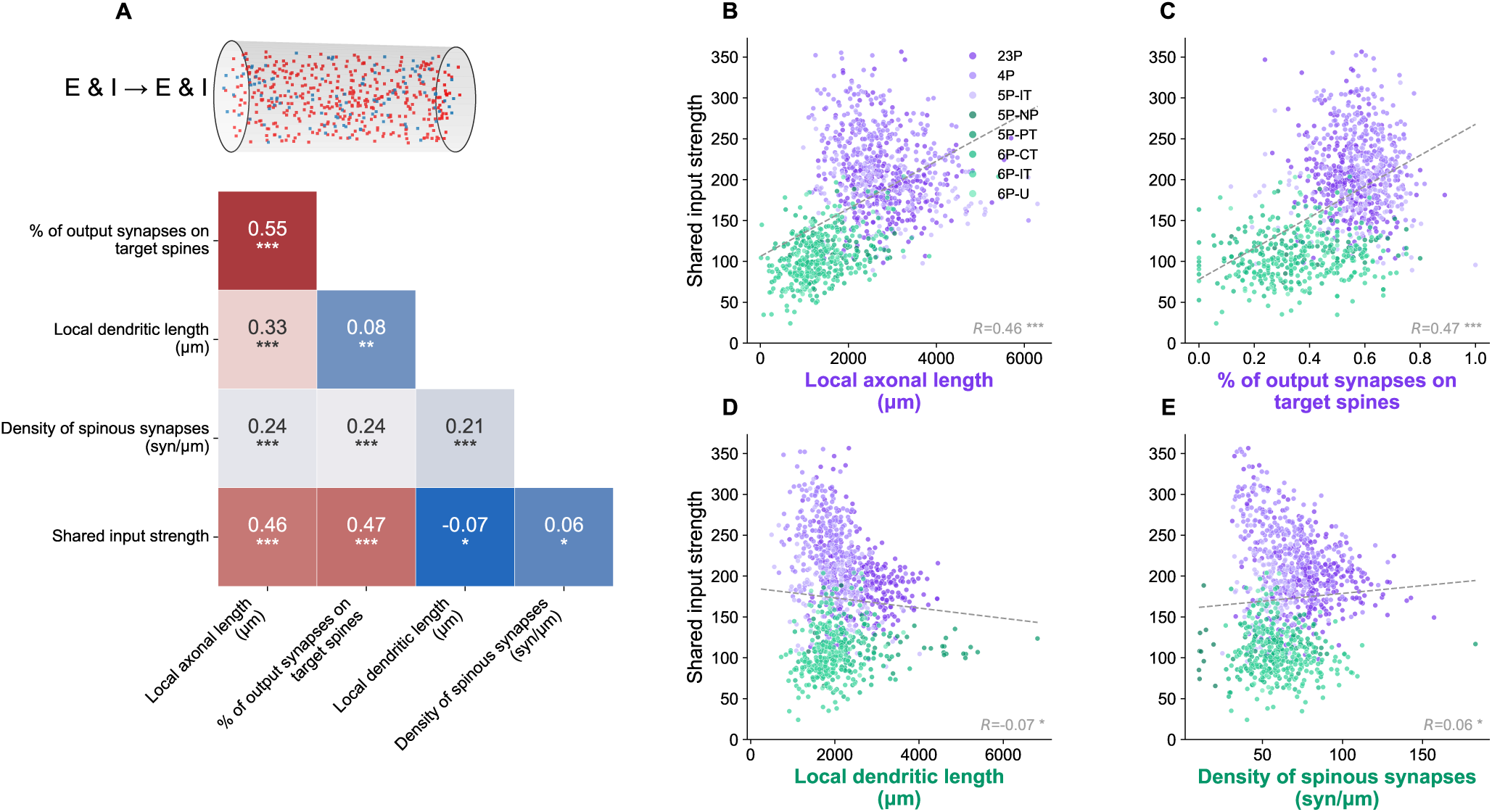
Feature correlations determining shared input strength across the full micro-column network (combined E and I inputs). This figure follows the exact same analysis and layout as Fig. S9, but extends the evaluation to the full network, incorporating both excitatory and inhibitory inputs to each neuron (matching the analysis conditions of main Fig. 4). **A:** A 5 × 5 lower-triangular Pearson correlation matrix showing the relationships between shared input strength and the four structural features for the full network. **B–E:** Full scatter plots of shared input strength (*y*-axis) versus the four individual features (*x*-axis), with high-common-input neurons indicated in purple and the rest in green. Similar to the isolated E-E network, “output” features B–C show stronger correlations (Pearson *R* = 0.46, *p <* 0.001 and *R* = 0.47, *p <* 0.001, respectively) with shared input strength than “input” features D–E (Pearson *R* = −0.07, *p* = 0.019 and *R* = 0.06, *p* = 0.045, respectively).

**Figure S11.**
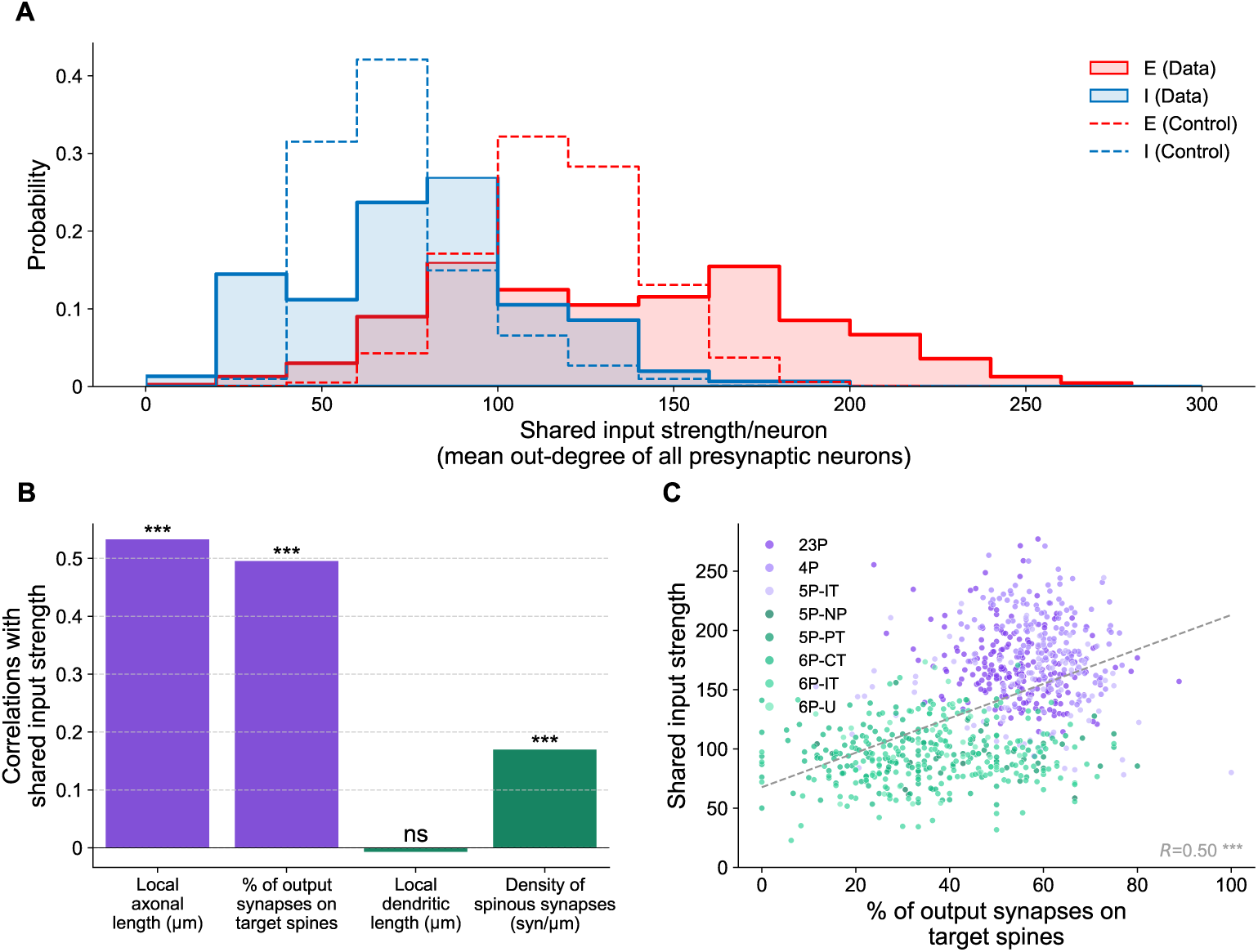
Shared-input architecture and preferential spine targeting excluding fully reconstructed axons. Analysis equivalent to Figure 4, performed on the “axon_partially_extended” neuronal subset (excluding 272 fully proofread excitatory micro-column neurons), where the input to each neuron consists of both excitatory and inhibitory connections. **A.** Distribution of the shared input strength per neuron for the excitatory (solid red) and inhibitory (solid blue) populations, compared against null distributions from 100 degree-preserving “blocked” configuration (CFG) models (dashed lines). **B.** Correlation of the shared input strength against four anatomical and connectivity features (purple: output features; green: input features). **C.** Scatter plot detailing the relationship between a neuron’s shared input strength and its spine preference (percentage of output synapses on target spines) across the entire micro-column population.

**Figure S12.**
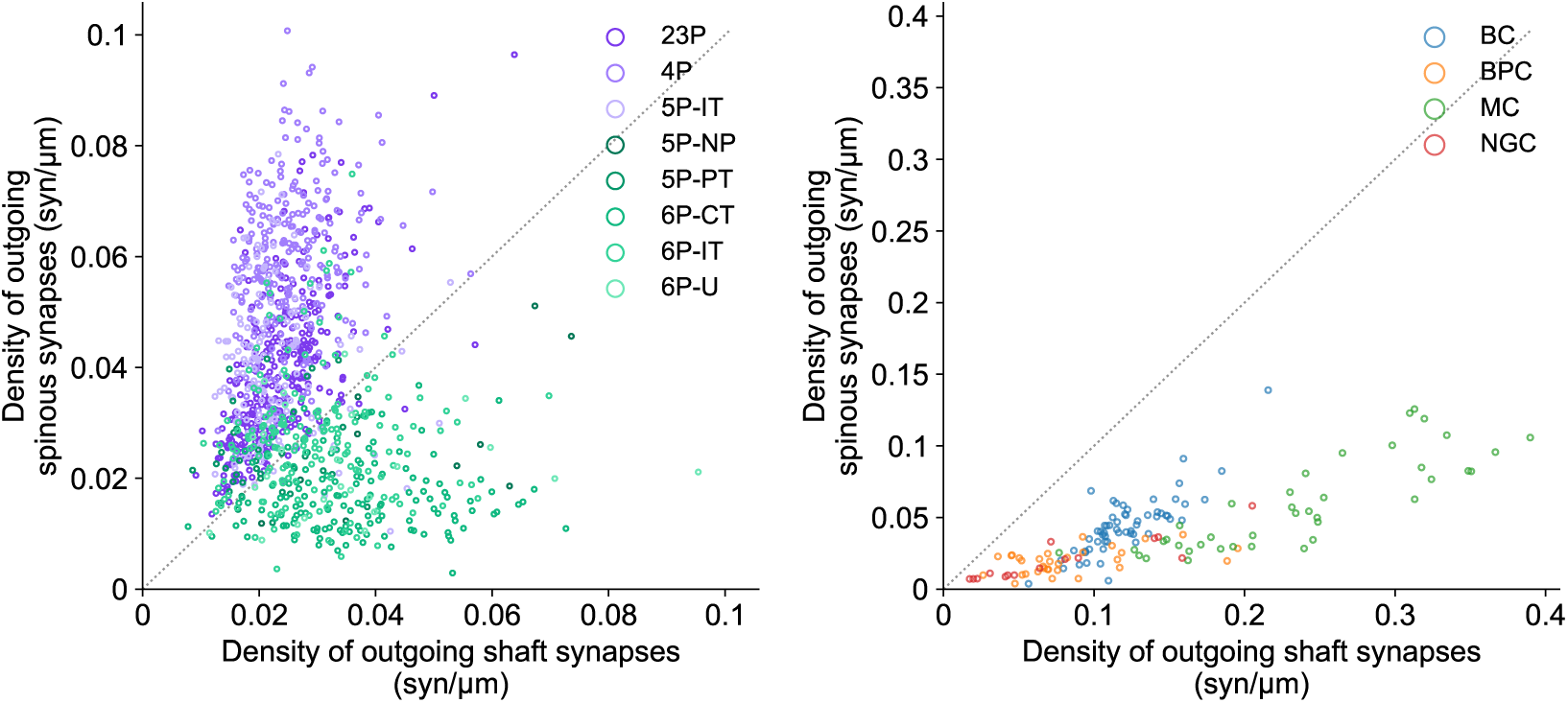
Subtype-specific axonal targeting strategies: Excitatory populations exhibit diverse target preferences, while inhibitory neurons uniformly target dendritic shafts. This is a whole-volume analysis evaluating the ∼1M outgoing synapses that project both within and outside the local micro-column. Direct comparison of outgoing spinous versus shaft synapse density for individual excitatory (Left) and inhibitory (Right) neurons, colored by cell subtype. The dotted grey line represents the 1:1 identity line (equal density on spines and shafts). *Left:* Excitatory neurons display broad, subtype-specific diversity in their targeting strategies. The high-sharing subpopulations (purple) predominantly target spines (clustering above the unity line), whereas the deeper-layer and output populations (green) exhibit a stronger preference for dendritic shafts (clustering below the unity line). *Right:* In contrast, all identified inhibitory subtypes exhibit a universal strategy, clustering entirely below the identity line to exclusively target dendritic shafts regardless of subtype classification.

**Figure S13.**
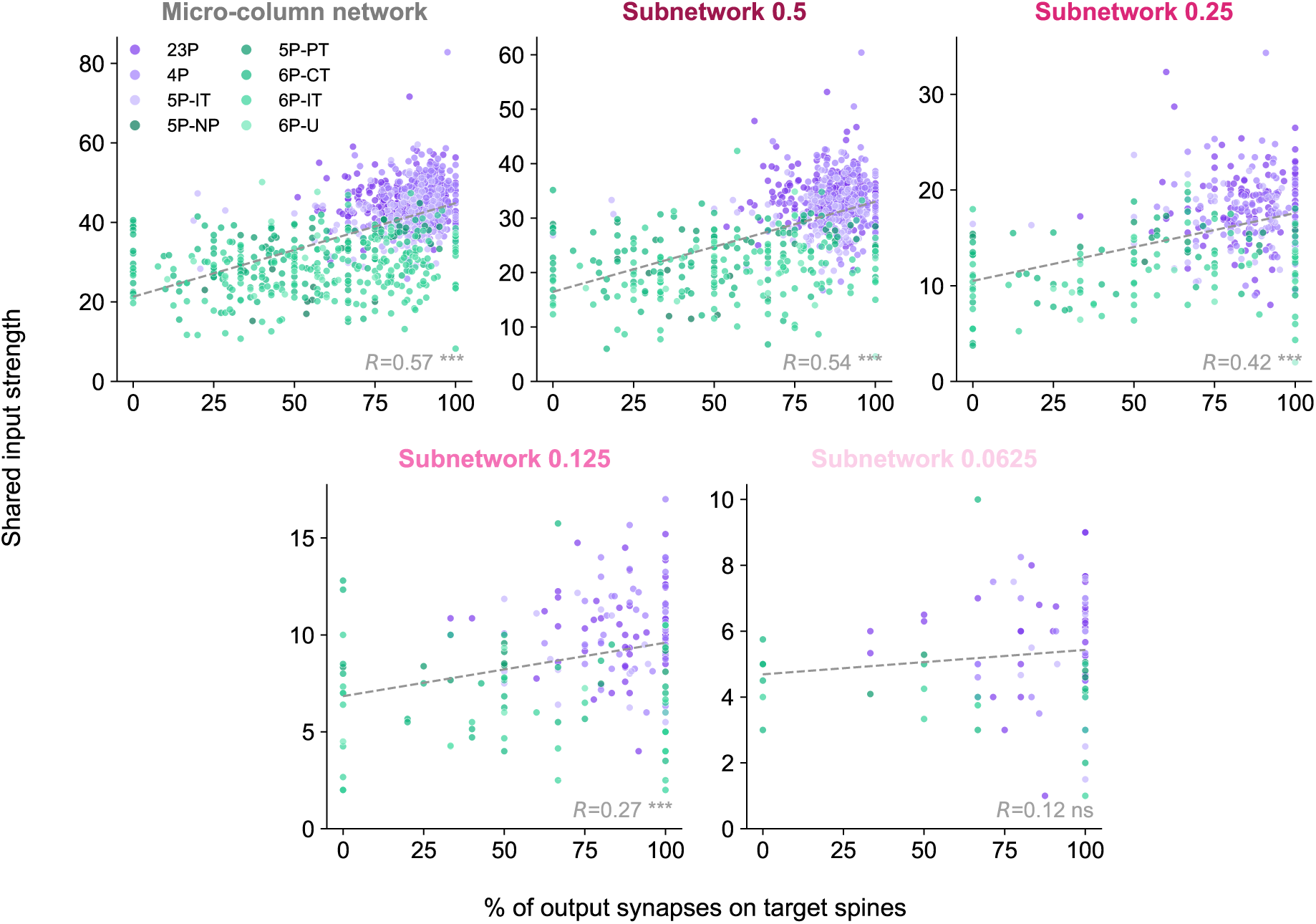
Correlation between shared input strength and percentage of output synapses on target spines across subnetwork scales. Corresponding to the analysis in Figure 5E. Neurons are colored by cell type: high-sharing, spine-preferring populations (purple) and deeper-layer, shaft-preferring populations (green). The positive correlations are maintained in larger volumes, but visible dispersion increases and correlation decays in the smallest subnetworks due to sparse sampling.

**Figure S14.**
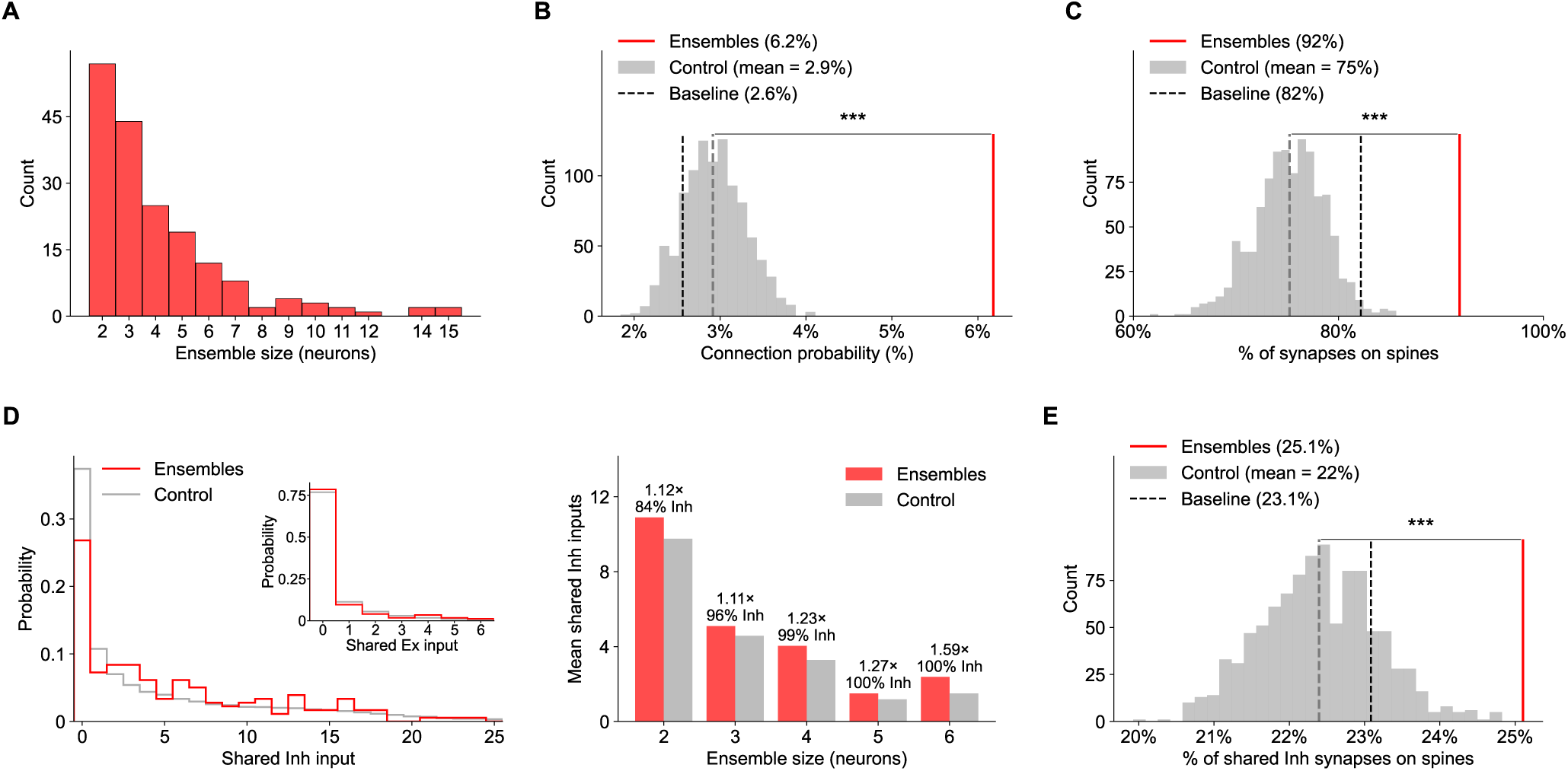
The wiring properties of coactive ensembles are independent of the ensemble-detection algorithm. All panels repeat the analyses of Figure 6 - identical metrics, matched controls, and paired bootstrap - on ensembles detected with a population-event clustering algorithm (Carrillo-Reid et al., 2015; Herzog et al., 2021; Ecker et al., 2024; see Methods) rather than the ICA-based detector. This yielded a larger and broader set of ensembles (*n* = 181, sizes 2–15), yet every result of Figure 6 is reproduced in direction, magnitude and significance. **A.** Distribution of ensemble sizes across all identified ensembles. **B.** Connection probability among ensemble members (red line; 6.2%) against a matched bootstrap null (grey; mean = 3%; 2.1-fold, *p* = 0.001), and the baseline E→E connection probability across the micro-column population (black dashed line; 2.6%). **C.** Percentage of the 209 synapses formed between ensemble members that target dendritic spines (red line; 92%) against a matched bootstrap null (grey; mean = 75%; *p* = 0.001), and the baseline E→E spine-targeting rate (black dashed line; 82%). **D.** Shared presynaptic input among ensemble members. *Left:* Distribution of the number of shared inhibitory presynaptic partners for ensembles (red) and matched controls (grey); 1,032 observed against a matched-null expectation of 875 (1.18-fold, *p* = 0.001). *Inset:* Same for shared excitatory partners; 128 against 85 (1.51-fold, *p* = 0.001). *Right:* Mean number of shared inhibitory partners by ensemble size, with fold-change and the inhibitory percentage of the ensembles’ shared input above each pair of bars. **E.** Percentage of shared inhibitory synapses onto ensembles’ dendritic spines (red line; 25.1%, over 9,986 synapses) against a null using the same matched controls as in D (grey; mean = 22%; *p* = 0.001), and the baseline I→E spine-targeting rate (black dashed line; 23.1%).

**Figure S15.**
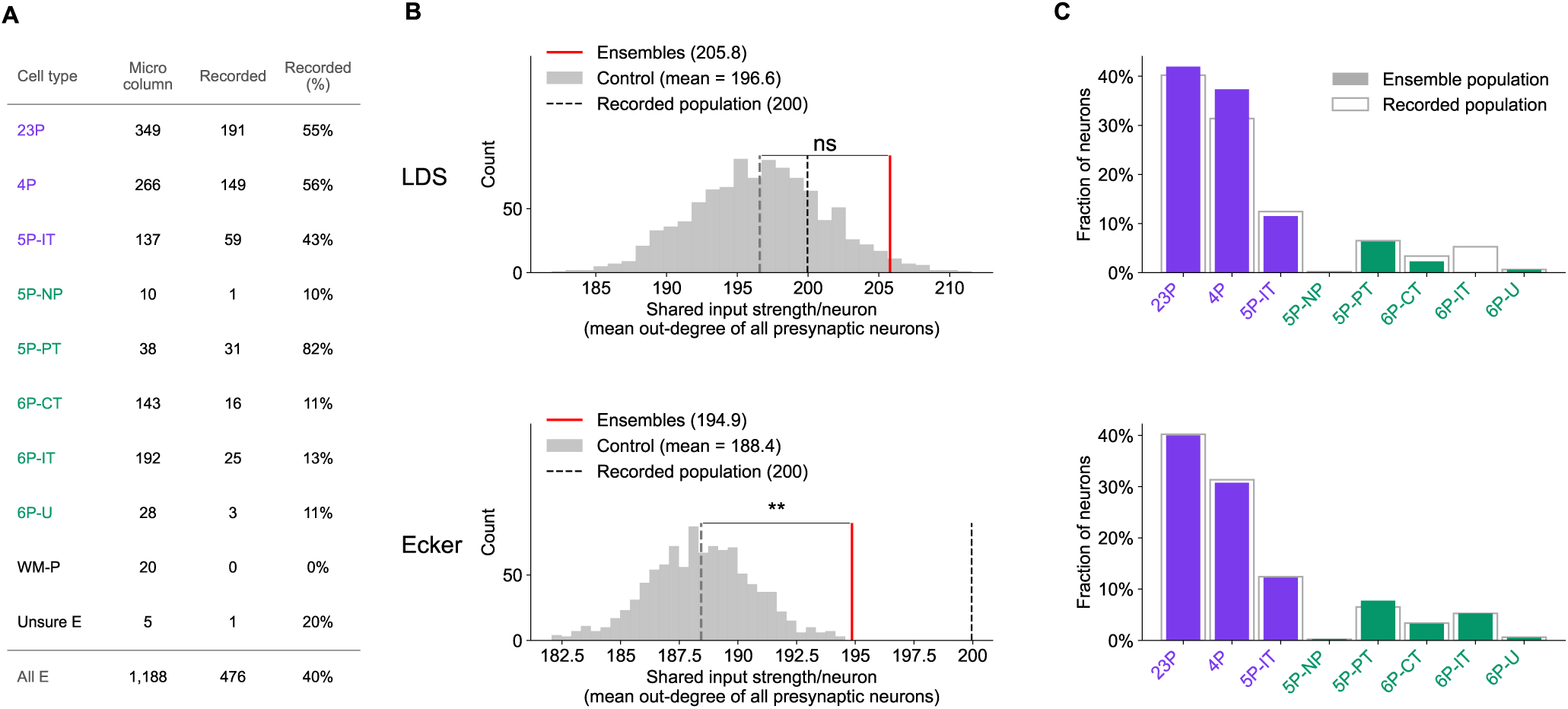
Ensemble members show a modest increase in shared input strength, consistent across both detection algorithms. Throughout the figure, cell-type names in the table (A) and the bars (C) follow the colour scheme of Figure 4F: purple marks the types with the highest shared input strength and spine-targeting preference (23P, 4P, 5P-IT), green the deep-layer and cortical-output types, which show lower values of both. **A.** Functional coverage of the micro-column: the number of excitatory neurons of each cell type in the micro-column, the number of those with calcium activity in at least one scan, and the resulting recorded fraction. In total, 476 of the 1,188 excitatory neurons of the micro-column (40%) were recorded; this pool is common to both detectors and is therefore the reference population used in C. The two unclassifiable types (WM-P, Unsure E) are listed here but omitted from C. **B.** Shared input strength of ensemble members - per neuron, the mean within-column out-degree of all its presynaptic partners (excitatory and inhibitory alike), the same measure plotted in Figure 4 - against the matched bootstrap null. The observed value averages over all *memberships*, i.e. every (neuron, ensemble) pair, so a neuron belonging to several ensembles contributes once per ensemble: 192 memberships for the first detector and 737 for the second, contributed by 174 and 401 distinct neurons respectively (C). *Top (LDS, Lopes-dos Santos* et al.*, 2013):* ensemble members (red line; 205.8) against matched controls (grey; mean = 196.6; 1.05-fold, *p* = 0.056, n.s.). *Bottom (Ecker, Ecker* et al.*, 2024):* ensemble members (red line; 194.9) against matched controls (grey; mean = 188.4; 1.03-fold, *p* = 0.002). Black dashed line, the mean over all recorded excitatory neurons (200). Each detector is compared only against its own matched control, so the two rows sit on different scales; expressed as the elevation of the members above their own control mean, the effect is in fact slightly smaller for the second detector (194.9 vs. 188.4, +3.4%) than for the first (205.8 vs. 196.6, +4.7%). It nevertheless reaches significance there because it is estimated over roughly four times as many memberships (737 against 192), i.e. the difference in *p* reflects statistical power rather than a stronger effect. **C.** Cell-type composition of the neurons appearing in at least one ensemble, counted once each (filled bars; *n* = 174 distinct neurons for LDS, *n* = 401 for Ecker), against the recorded excitatory population (open outline), for each detector. Both compositions broadly track the recorded population: LDS is enriched for 4P (1.19-fold, *z* = 2.2) and returns no 6P-IT members (0 observed against 9.1 expected, *z* = −3.9), while Ecker matches the recorded population across all types apart from a 5P-PT enrichment (1.19-fold, *z* = 2.5).

## Footnotes

1 https://tutorial.microns-explorer.org/annotation-tables.html

2 https://tutorial.microns-explorer.org/proofreading.html

3 Software implementation provided by R. Urlus; https://github.com/RUrlus/diptest

4 https://github.com/cajal/microns_phase3_nda

